# Noninvasive Sleep Scoring in Mice using Electric Field Sensors

**DOI:** 10.1101/794552

**Authors:** H Kloefkorn, LM Aiani, A Lakhani, S Nagesh, A Moss, W Goolsby, JM Rehg, NP Pedersen, S Hochman

**Affiliations:** Department of Physiology, School of Medicine, Emory University, Atlanta, GA, USA; Department of Neurology, School of Medicine, Emory University, Atlanta, GA, USA; School of Electrical and Computer Engineering, Georgia Institute of Technology, Atlanta, GA, USA; School of Interactive Computing, Georgia Institute of Technology, Atlanta, GA, USA

**Author notes:** Corresponding Authors’ full addressed and current emails: Heidi Kloefkorn, 615 Michael Street, Department of Physiology, Atlanta, Georgia, 30033, Nigel P. Pedersen, 101 Woodruff Circle, Department of Neurology, Atlanta, Georgia, 30322.

**Keywords:** 3-state Sleep, Sleep Scoring, Electric Field Sensor, Noninvasive, Rodent

## Abstract

**Background:** Rodent sleep scoring in principally reliant on electroencephalogram (EEG) and electromyogram (EMG), but this approach is invasive, can be expensive, and requires expertise and specialized equipment. Affordable, simple to use, and noninvasive ways to accurately quantify rodent sleep are needed.

**New method:** We developed and validated a new method for sleep-wake staging in mice using cost-effective, noninvasive electric field (EF) sensors that detect respiration and other movements. We validated recordings from EF sensors attached to the exterior of specialty chambers used to continuously capture sleep with EEG/EMG, then compared this to EF sensors attached to vivarium home-cages.

**Results:** EF sensors quantified 3-state sleep architecture (wake, rapid eye movement – REM – sleep, and non-REM sleep) with high agreement (>93%) and comparable inter- and intra-scorer error as expert EEG/EMG scoring. Novices given an instruction document with examples were able to score sleep comparable to expert scorers (>91% agreement). Additionally, EF sensors were able to quantify 3-state sleep scoring in traditional mouse home cages.

**Comparison with existing method:** Most noninvasive sleep assessment technology requires animal contact, altered cage environments, and/or can only discern 2 states of arousal (wake or asleep). The EF sensors are able to discriminate REM from non-REM sleep accurately and from outside the animal’s home cage.

**Conclusions:** EF sensors provide a simple and reliable method to accurately score 3-state sleep architecture; (i) from outside the typical home cage, (ii) where noninvasive approaches are preferred, or (iii) which EEG/EMG is not possible.

**Graphical Abstract:** 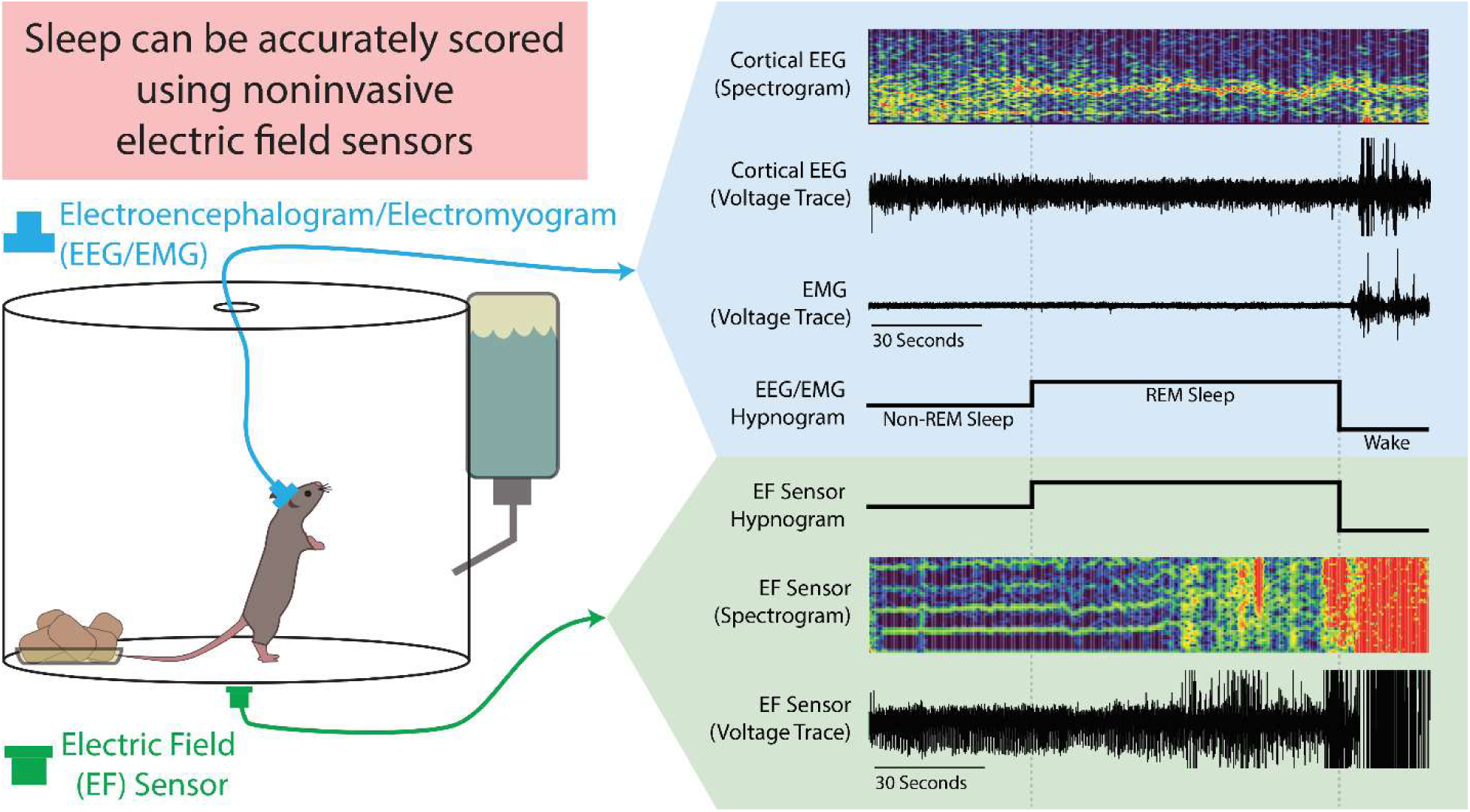

## Introduction

Accurately characterizing sleep-wake is crucial to understanding its impact on health, cognition, and injury recovery.^1–3^ Sleep staging in humans, relies on substantial non-invasive instrumentation (polysomnography) including electroencephalogram (EEG), electromyogram (EMG) of chin and limbs, electrooculogram (EOG) to detect rapid and rolling eye movements, along with chest wall or air flow movement, oximetry, and video.^4–6^ However, rodent studies typically rely on fewer signals, invasive implanted EEG and EMG, less commonly invasive EOG electrodes, and video analysis. Invasive surgical implants can result in weight loss, extensive recovery, and commonly requires use of a tether cable.^7–9^ Moreover, surgical expertise, limited locations for electrode placement, and specialized equipment required to collect EEG and EMG data may restrict inclusion of sleep analysis in an experimental design. For these reasons, there is a need to develop noninvasive methods to assess sleep in preclinical rodent models.

Most noninvasive rodent sleep assessment methods center around measures of gross body movement. Several video analysis techniques have been used to distinguish between 2-state conditions (wake or asleep), but are unable to reliably define 3-state sleep architecture that further differentiates sleep into rapid-eye-movement (REM) and non-REM sleep.^10–12^ In individually housed animals, other remote methods using light reflection and Doppler techniques have been shown to distinguish REM from non-REM sleep in rats,^13^ but not in mice.^14^ Alternatively, force sensors placed inside modified rodent cages have used gross body movement to quantify sleep.^7,15–18^ These force sensors require animal contact, individual housing, and special analytical software to characterize sleep, but only one study was able to measure 3-state sleep architecture in mice.^7^

Respiration is a distinguishing factor for successfully measuring 3-state sleep architecture through noninvasive movement-based measures, as respiration profiles can distinguish REM from non-REM sleep architecture in humans,^5,19^ dogs,^20,21^ and in rodents.^8,22^ The more successful noninvasive approaches to quantify sleep incorporate respiration-related measures into their models^7,8,13,15,17^ and one study was able to reliably determine 3-state sleep architecture in mice through respiration alone using whole body plethysmography.^8^

In this study, a new method to quantify 3-state sleep architecture was developed using electric field (EF) sensors. EF sensors continuously detect fluctuations in the local static electric field caused by motion and have been shown to capture both gross body movement and respiration-related measures.^23^ EF sensors were mounted on the outside surface of cylindrical acrylic chambers housing mice instrumented with EEG and EMG electrodes. Simultaneously-collected data from EEG, EMG, and EF sensors were used to create rules for scoring sleep using EF sensors alone. A data set was recorded and then divided into separate EEG/EMG and EF recordings, scored by experts, then compared to determine whether EF sensors could reliably and accurately quantify 3-state sleep architecture. Additionally, novices were recruited to assess ability to accurately score sleep using EF scoring alone. Overall, EF sensor data alone can reliably and easily distinguish 3-state sleep architecture with comparable precision and accuracy as EEG/EMG and that novices can learn how to accurately score sleep from EF sensor data alone.

## Methods

### Animals

All work was conducted according to FASEB and Emory IACUC animal guidelines. Four adult mice (3-6 months old: wildtype female n=1, Vgat^flox^ female n=1, male n=2, all backcrossed to C57BL/6J), participating as controls in another study, were used to validate the EF sensor technology.

### Implanting EEG/EMG Electrodes

Hippocampal and contralateral fronto-parietal cortical EEG and EMG were recorded using custom-made headsets described in another publication.^24^ Briefly, mice were deeply anesthetized and placed on a stereotaxic frame. Meloxicam (5mg/kg, 1cc) was administered subcutaneously (intrascapular). The skull was prepared for aseptic surgery and lidocaine (2% in saline) was administered locally. The skull was exposed using a scalpel and surgical scissors, cleaned, and scored with the reverse side of a scalpel blade to provide a better substrate for dental cement adhesion. To accommodate EMG electrodes, the skin over the neck was blunt-dissected away from muscle. The mouse head was then leveled within the stereotaxic frame and holes were drilled stereotaxically to match the corresponding location of electrodes on the headplate. The headplate was then lowered into place and attached via screw electrodes, cyanoacrilic adhesive, and dental cement. EMG electrodes were placed beneath the prepared skin, but above the muscles. The wound was closed by drawing the skin over the adhesives and suturing (5-0 silk suture). The animals recovered for four days prior to tethering the EEG/EMG headplate to a preamplifier (Pinnacle Technology, Inc., 8406-SE31M, 100x gain), communicator (Pinnacle Technology, Inc., 8408), and A/D converter (Power1401-3A, Cambridge Electronic Design, UK). Recordings used for this study were collected more than one month after surgery.

### Electric Field Sensors

Electric field (EF) sensors (Plessey Semiconductors, PS25251, 1 cm2, +/- 5V) were adapted to interface with an A/D converter (Power1401-3A, Cambridge Electronic Design, UK). These EF sensors measure changes in the local electric field caused by movement and translate it into a voltage trace (+/-5V, 2048Hz sampling rate). While EF sensors have a frequency response range within 0.1hz −10 kHz, a two-pole low pass filter set at 12 Hz was used to exclude interference from higher frequency events. Recording features and validation of these EF sensors to measure respiration in rodents have been published previously.^23^ Briefly, the EF sensors can measure animal motion with high resolution from outside the home-cage. Moreover, these EF sensors are able to reliably measure extremely small mouse motions with great sensitivity, allowing them to detect respiratory-related movement, validated against whole body plethysmography, when the animal is at rest.^23^

During Non-REM sleep, the predominant animal movement is caused by respiration via chest wall expansion and contraction. During REM sleep, respiration depth and frequency vary in addition to twitches of the distal digits of the limbs, tail, whiskers, and ears.^25^ In this study, these state-related movement changes were detected by EF sensors and validated against EEG and EMG.

### EEG/EMG and EF Recording Set-up

To accommodate EEG/EMG recording, animals were housed singly in cylindrical acrylic chambers (130 oz. clear acrylic container, Oggi Corporation, Anaheim, CA) with corncob bedding, nesting material, and free access to water and food. The lid contained a hole appropriate for the tether associated with the EEG/EMG recording equipment (Figure 1). Cameras (1 megapixel day/night dome USB security camera, ELP, 320, 30 frames per second, Shenzen, China) were placed above each cage to synchronously visualize the cage environment. Two EF sensors were placed on the exterior bottom of each cylindrical chamber; only one was needed for characterization of respiratory changes during sleep. Housing room humidity was kept between 30-70%. Recordings were captured continuously for one week, in 12 hour segments, using Spike2 software and automated acquisition (Spike2 v8.09, CED, UK, 2048Hz sampling) with synchronization of the video, EEG, EMG, and EF signals. Although EF sensor data were collected here using CED hardware and Spike2 software, they can be recorded and analyzed using any other equipment that permits analog signal acquisition, such as Axon instruments Digidata/pCLAMP (Molecular Devices), National Instruments/LABVIEW, or MATLAB.

**Figure 1.**
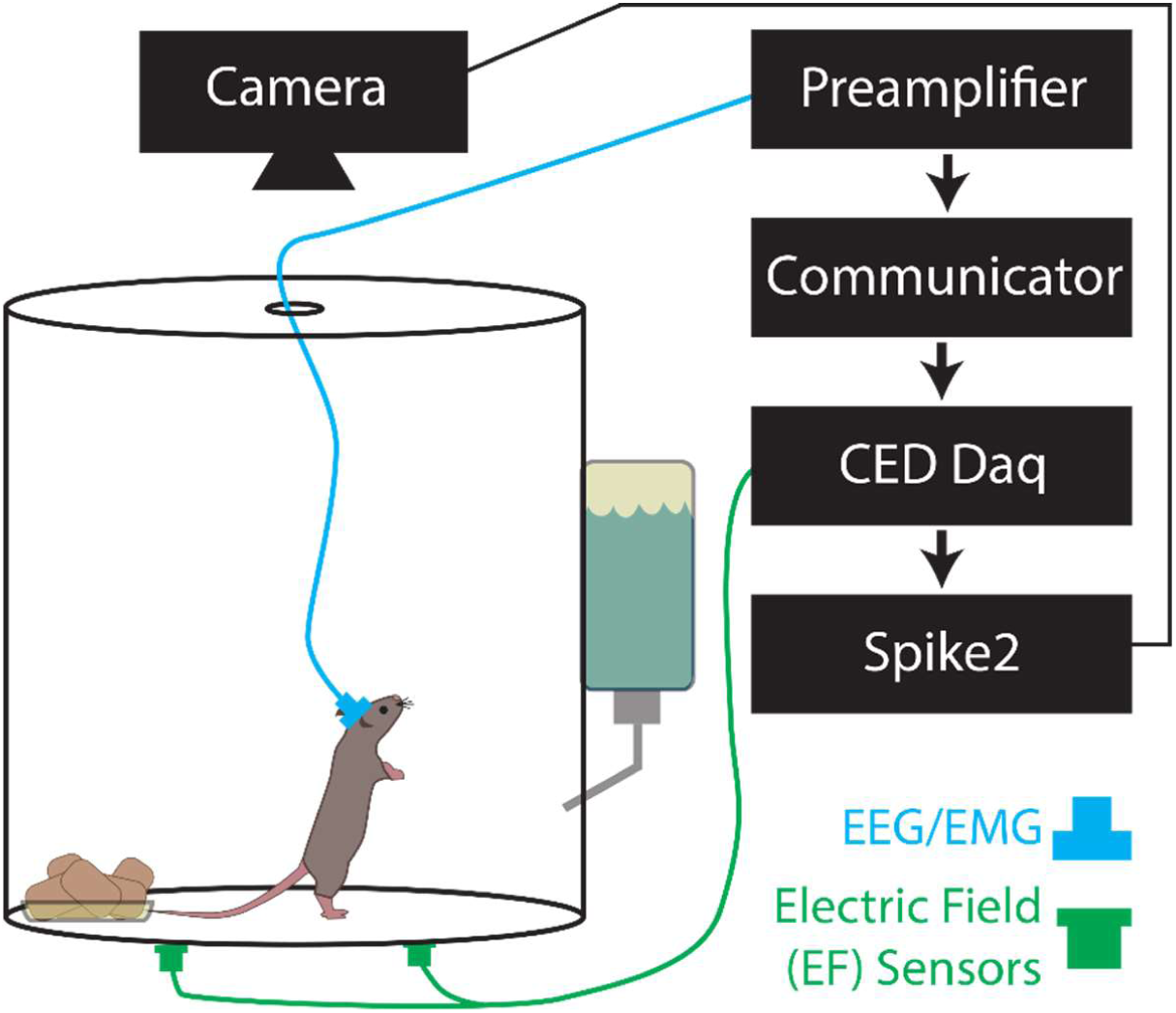
Cage Set-up for Electroencephalogram/Electromyogram (EEG/EMG) and Synced Electric Field (EF) Recordings. The animals were singly housed in cylindrical acrylic chambers. Head stages (blue) were surgically mounted prior to recordings and contain EEG and EMG electrodes connected to a preamplifier, communicator, then a Cambridge data acquisition box (CED Daq). EF sensors (green) were attached to the cage exterior and also connected directly to the CED Daq. A camera was mounted above the cage to visually record animal activity but was not used for sleep scoring.

### Scoring Sleep

Two 5.5 hour recording samples in four mice (totaling 8 recordings), randomly distributed within the 24-hour day period, were chosen to score sleep. Features and descriptions of each recording are summarized in Table 1. This design, with shorter than 24-hour epochs was intended to test whether EF sensor scoring could accurately discriminate 3-state sleep scores by comparison to conventional scoring.

**Table 1.** Description of recordings used to validate electric field (EF) sensors. Recordings are taken from 4 animals at all times during the circadian cycle specifically to assess whether EF sensors can accurately quantify 3-state sleep architecture. The purpose of this design is not to describe the sleep-wake behavior of these animals over a 24 hour period. Each recording consisted of 5.5 hours of data and was scored by 3 experts using electroencephalogram (EEG) and electromyogram (EMG) to determine 3-state sleep architecture and the resulting percentage of each arousal state for each recording is given.

|  | Recording<br>1 | Recording<br>2 | Recording<br>3 | Recording<br>4 | Recording<br>5 | Recording<br>6 | Recording<br>7 | Recording<br>8 |
| --- | --- | --- | --- | --- | --- | --- | --- | --- |
| Animal | Animal 1 | Animal 1 | Animal 2 | Animal 2 | Animal 3 | Animal 3 | Animal 4 | Animal 4 |
| Time of Day (24-hour) | 16:00 | 8:00 | 11:00 | 13:00 | 22:00 | 4:00 | 8:00 | 2:00 |
| Total Number of 10-second Epochs | 1572 | 1643 | 1977 | 1345 | 1977 | 1977 | 1977 | 1977 |
| EEG/EMG Scored as wake (%) | 34.9 | 37.2 | 43.2 | 56.1 | 60.1 | 35.8 | 37.6 | 17.0 |
| EEG/EMG Scored as non-REM sleep (%) | 54.6 | 54.4 | 50.4 | 37.4 | 33.1 | 59.5 | 53.1 | 71.2 |
| EEG/EMG Scored as REM sleep (%) | 10.5 | 8.4 | 6.4 | 6.5 | 6.8 | 4.7 | 9.3 | 11.8 |

For each of the 8 recordings prepared for the development and validation of scoring, channels with EF signals and EEG/EMG were split to enable blinding to the other signal (Figure 2A). In this way, each recording was scored purely on the basis of EEG/EMG or EF data without needing the video data. Recordings were then presented in random order and scored blindly; scorers knew whether the file contained EEG/EMG or EF data because different scoring rules were required for each method, but they did not know which EEG/EMG derivative recording went with its twin EF derivative recording. In addition, to assess intra-scorer reproducibility, three recordings each of EEG/EMG and EF taken from the same original recording were randomly chosen and blindly scored again within the data set.

**Figure 2.**
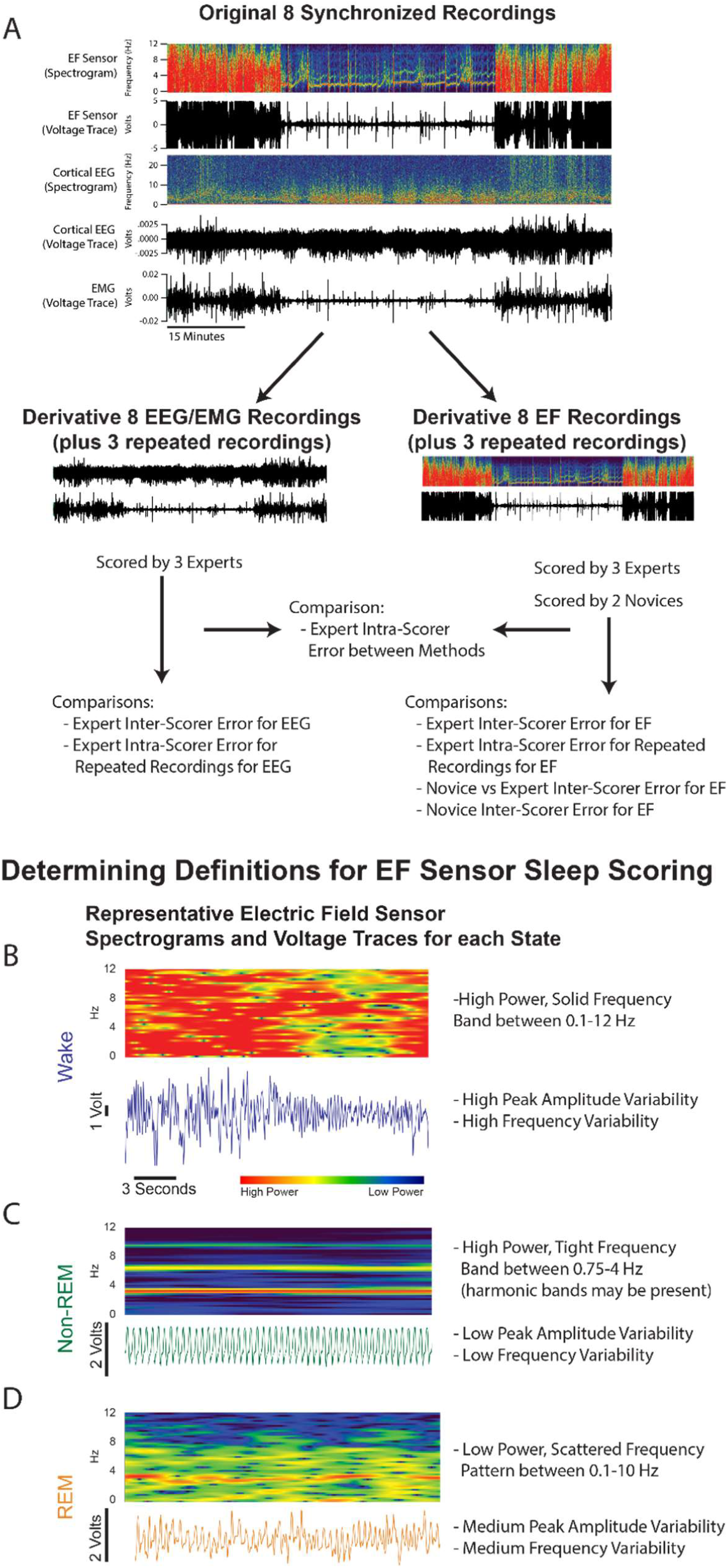
Recording Set-Up for and Creating Definitions for Scoring. A) Each recording is created as synchronous voltage traces from electric field (EF), electroencephalogram (EEG), and electromyogram (EMG). As an example, one EF and EEG voltage trace are also represented as a spectrogram – a graphic in which the x-axis is time, the y-axis is frequency, and the color intensity denotes the power of respective frequencies present in the voltage trace. Each synchronized recording is divided into two files that contain either the EEG/EMG or EF voltage traces for subsequent scoring and comparison. B) The EF sensor wake state data appears as both a spectrogram (top) and voltage trace (bottom in navy). Each voltage trace is the direct output from the EF sensors and represent animal movement. C) EF data for non-REM sleep behavior. D) EF data for REM sleep behavior.

### Defining Criteria for Scoring Sleep using EF Sensors

All recordings were scored manually using Spike2 by three expert sleep scorers. For the EEG/EMG recordings, sleep state (wake, REM sleep, or non-REM sleep) was manually scored in accordance with conventional Rechtschaffen and Kales^26^ criteria in 10-second epochs using only the cortical EEG electrode and EMG. Expert sleep scorers, with extensive experience scoring sleep-wake in mice, used 12 hours of EEG/EMG scored behavior to develop criteria for identification of sleep state by EF sensor recordings. EF sensors were found to reliably detect movement-related features that can discriminate between wake, non-REM, and REM states based on frequency domain analysis using Fast Fourier transform (FFT) as show in Figure 2 and detailed below. A reference instruction document was generated to score sleep exclusively by EF sensor criteria (Supplement 1).

During wake, the raw voltage output of the EF sensors (Figure 2B) is of high amplitude and erratic, reflecting the highly complex summations of movement for motor activity. Wake-state frequency domain patterns on FFT generally include multiple signals between 0.1-12 Hz and amplitude generally ranges 4-10 V (as measured between ±5V). These raw voltage traces were converted into Spike2 ‘sonograms’ which are FFT spectrogram plots (Hanning filter, 2 second overlapping bins) characterizing the frequency signals (y-axis, Hz) across time (x-axis, seconds) and their relative power (color intensity) that make up the raw voltage trace. During wake, the spectrogram shows intensely powerful (i.e. red) frequencies that solidly span from near 0 Hz to 12 Hz.

During non-REM sleep, the raw voltage output of the EF sensors is consistently rhythmic and correlated with the repetitive motion of breathing during Non-REM sleep (Figure 2C). Non-REM-state frequency patterns generally includes a single signal between 2-4 Hz and amplitude generally ranges 0.1-2 V. A non-REM spectrogram, likewise, exhibits a powerful (i.e. red), consistent frequency band associated with respiratory rate typically occurring between 2-4 Hz. Occasionally, less powerful harmonic artifacts at whole number multiples of respiration frequency will appear in the spectrogram of non-REM sleep given the non-sinusoidal nature of the signal.

During REM sleep, the raw voltage output of the EF sensors increases in frequency variability relative to non-REM, but amplitude changes are minimal (Figure 2D). REM-state frequency patterns generally include multiple signals from 1-10 Hz and amplitude generally ranges 0.4-3 V. Spectrograms of REM sleep show these multiple frequency signals as fragmented spots of frequencies ranging from just below the corresponding non-REM respiratory rate (2-4 Hz) to 10 Hz. The frequency band associated with respiration often appears on the spectrogram during a REM event, but becomes less consistent in both frequency and intensity. The relative power of these cumulative signals during REM sleep is typically equivalent or slightly less than the corresponding non-REM power, but much less powerful than during wake. Overall, as shown in Figure 2, REM and non-REM states are easily discriminable.

### Scoring Sleep using EF Sensors

Three expert sleep scorers used the developed criteria for scoring sleep using EF sensors. Additionally, two novice scorers used the same criteria to score the same EF recordings under two different conditions: Novice #1 scored EF recordings given only the EF sleep scoring training document (Supplement 1) while Novice #2 was also given two example recordings of sleep-scored EF sensor data to aid assessment of scoring (Supplement 2).

### Statistical Validation of EF Sensor Sleep Scores to EEG/EMG Sleep Scores

EEG/EMG and EF recordings were scored independently by the three expert sleep scorers. Comparisons were made on an epoch-by-epoch basis. Intra-scorer (e.g. Scorer #1 EF compared to Scorer #1 EEG/EMG) and inter-scorer (e.g. scorer #1 EF compared to Scorer #2 EF) percentage agreement was calculated from the total number of epochs and represents consensus agreement – percent of epochs in which all 3 expert scorers agreed. 2-way paired t-tests with an alpha of 0.05 were performed on the percentage agreement measures.

To determine the reliability and reproducibility of EF sensor scoring and EEG/EMG scoring, intra-class correlation coefficients (ICCs) were calculated. ICCs describe how well data scored across graders or methods resemble each other accounting for grouping rather than only as paired observations. Where appropriate, single measure ICCs were reported for intra-scorer comparisons, average measures ICCs were reported for inter-scorer comparisons, and 2-way paired t-tests with an alpha of 0.05 were calculated to compare ICCs between scoring methods.

The two novice EF scores were compared to each of the three expert scores individually to produce three different percentage agreements and ICCs per novice. These percentage agreements and ICCs were averaged and compared using 2-way paired t-tests with an alpha of 0.05.

Percentage agreement and descriptive correlation statistics (ICCs, Pearson’s R, or Cohen’s kappa) are the most commonly reported variables to compare two sleep scoring methods. They are representative of how well broad sleep measures might perform (i.e. total time asleep), but not other measures that require more detail (e.g. sleep fragmentation). To assess how EF sleep scoring matches EEG/EMG sleep scoring with more resolution, the number of transitions between arousal states (into wake, into non-REM sleep, and into REM sleep) were calculated for each scorer and recording. Differences between state transition values for EF and EEG/EMG methods were assessed using 2-way paired t-tests with an alpha of 0.05. Unless otherwise stated, data are presented as mean ± standard deviation.

Sensitivity (false positivity rate) and specificity (false negative rate) were calculated for intra-scorer agreements (e.g. Scorer #1 EF compared to Scorer #1 EEG/EMG, where EEG/EMG is considered the ground truth) and novice agreements with expert scorers (where the expert score was considered the ground truth). Sensitivity and specificity are also reported broken down by arousal state (wake, non-REM sleep, and REM sleep).

### Applying EF Sensors to Traditional Mouse Home Cages to Assess Sleep

Once the EF sensor technology was validated against EEG/EMG in the cylindrical chambers to accurately assess sleep, three typical rectangular vivarium mouse home-cages were instrumented with EF sensors to determine whether sleep could be measured in an environment without EEG/EMG methods. In these experiments, 6 female C57BL/6 mice (3 months old) were pair-housed in standard 12:12 hour light, food, water, and temperature home-cage conditions. During testing, the home-cages were temporarily divided for the 12 hour dark cycle into two electrically shielded compartments in which the mice were still able to see, smell, and hear their cage-mate. EF sensors were attached to the home-cage exterior allowing each animal to be recorded individually (Figure 3A). Once acclimated to the divided cage set-up, data were collected for 12 hours during the dark cycle using the Digidata/pClamp acquisition platform (Molecular Devices). Collected data was then imported into Spike2 (CED) where sleep was scored by one of the expert scorers.

**Figure 3.**
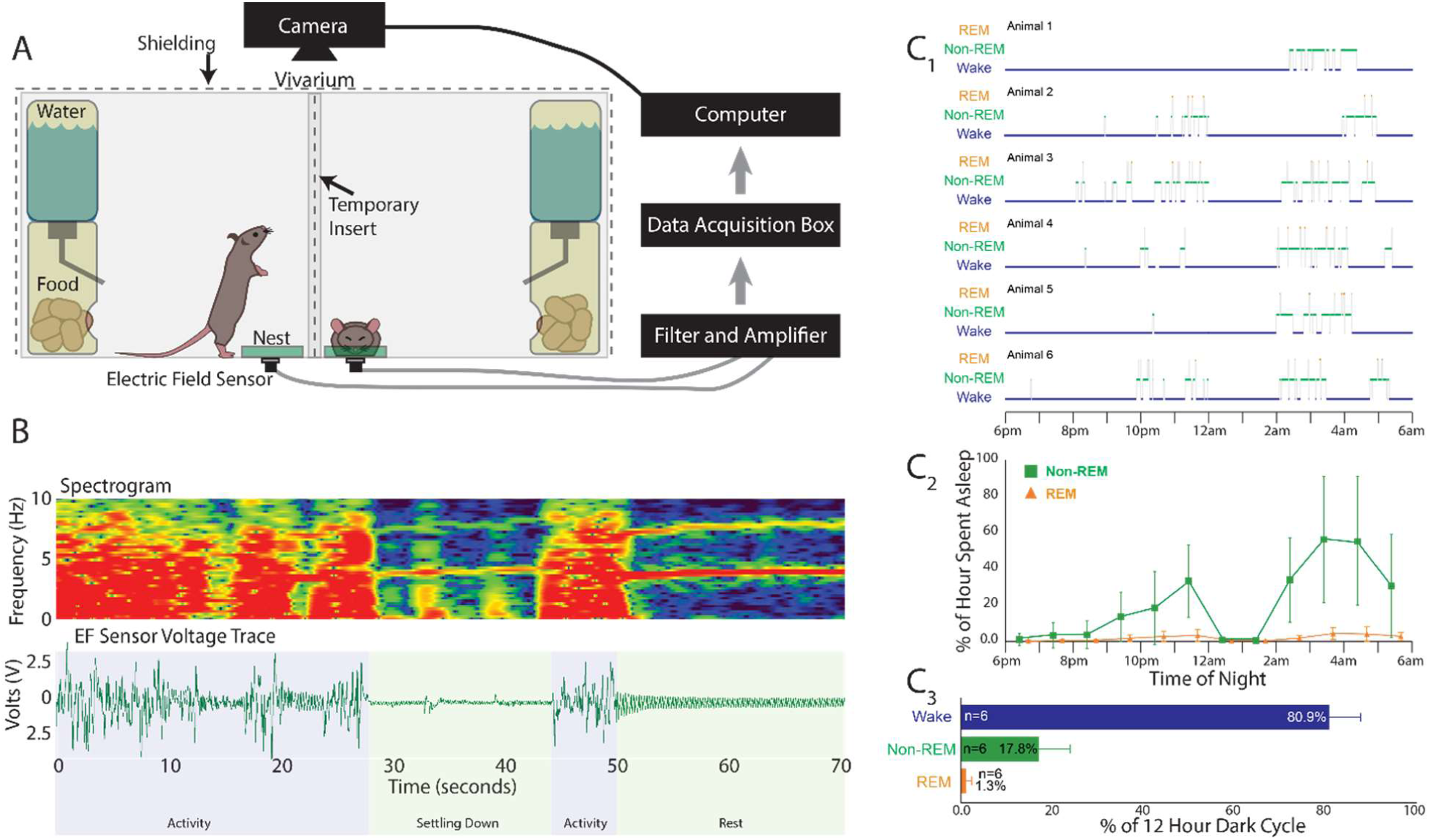
Description of the Electric Field (EF) Sensors Applied to Animal Home Cages without Electroencephalogram/Electromyogram (EEG/EMG) Validation. A) Home cage set-up in which the animals are separated by a transparent shielded insert during testing that allows visual, olfactory, and thermal interactions between the animals. Each animal has free access food and water and a 60 mm petri dish to use as a nest. The electric (EF) sensors are attached to the home cage exterior and connected to a filter/amplifier box, an Axon instruments data acquisition box, then to the computer. B) Representative spectrogram and raw voltage trace from home cages that are indistinguishable from the data collected on the electroencephalogram/electromyogram (EEG/EMG) validation cages. C_1_) 3-state hypnogram describing the sleep results for 6 animals recorded overnight between 6pm and 6am. C_2_) The average percentage of sleep (non-rapid eye movement – non-REM – sleep plus REM sleep time) of all 6 animals per hour over the 12 hours. C_3_) The 12-hour average of all 6 animals for each arousal state. Data are presented as mean ± standard deviation.

## Results

### Sleep scoring had high agreement within and between EEG/EMG and EF scoring methods

The three expert scorers were compared within each sleep scoring method, EEG/EMG or EF, to determine method-related error. Within both EEG/EMG and EF scoring methods, consensus agreement of the expert scores were above 93%, comparable with prior reports,^27^ and ICCs were above 0.97 (Table2; Fig4A).

**Table 2.**
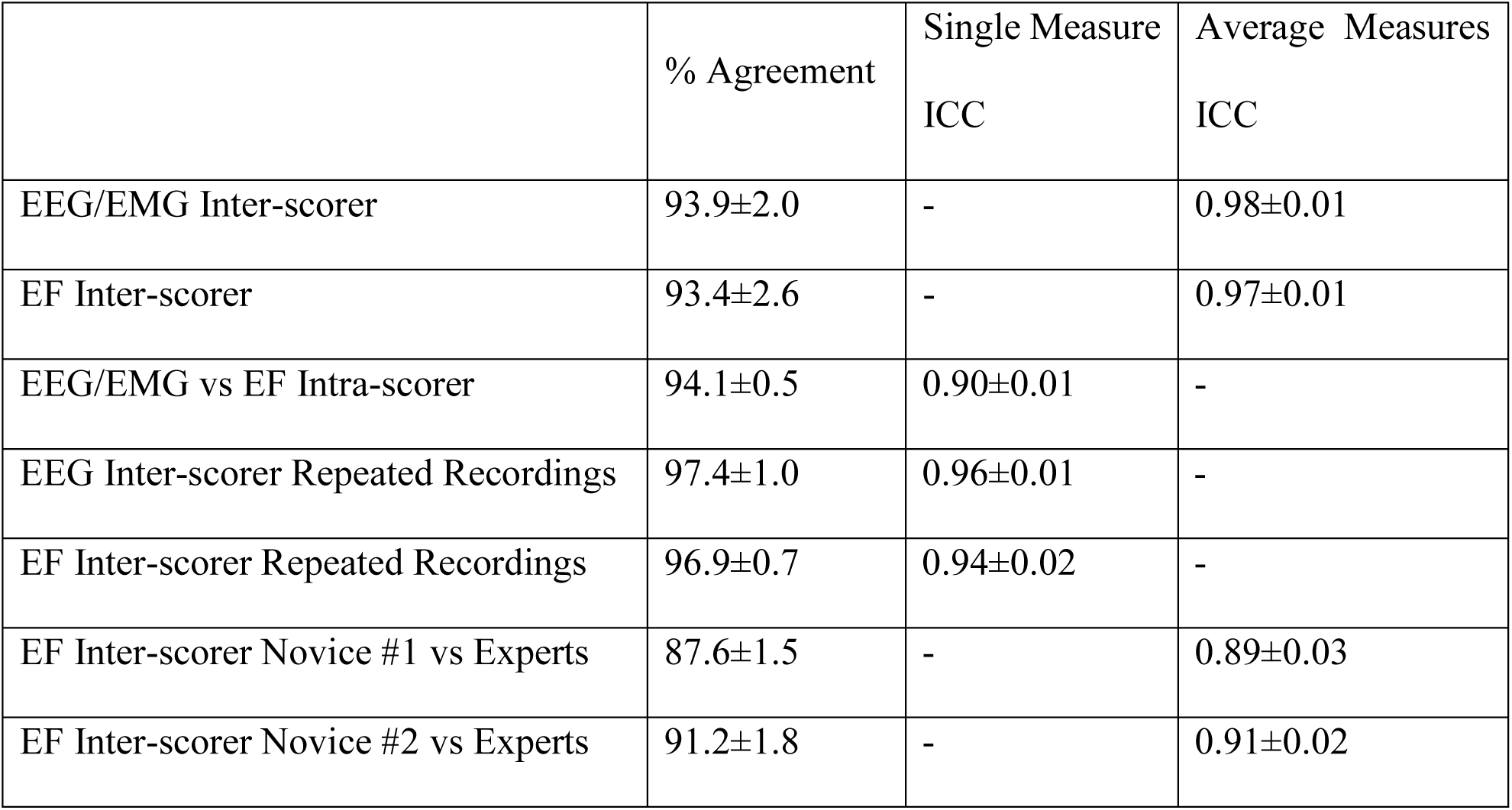
Summary of % agreement and intra-class correlation coefficients (ICCs) for different comparisons in validating electric field (EF) sensors sleep performance against the electroencephalogram (EEG) and electromyogram (EMG) method. Data presented as mean ± standard deviation.

**Figure 4.**
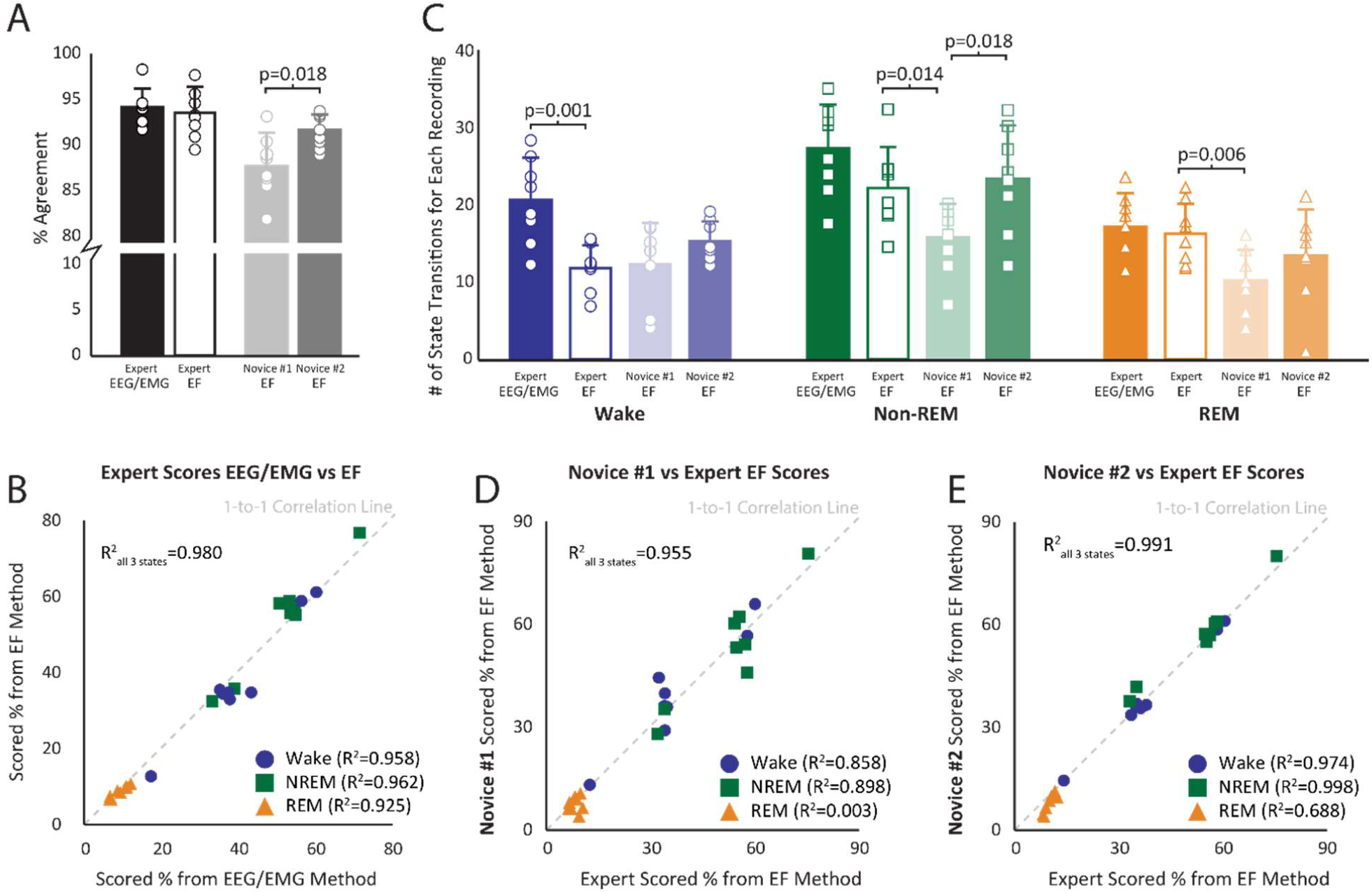
Agreement Between Electric Field (EF) and Electroencephalogram/Electromyogram (EEG/EMG) Sleep Scoring Methods. A) The black and white bars represent expert intra-scorer agreement for both electric field (EF) and electroencephalogram/electromyogram (EEG/EMG) methods. The light and dark gray bars represent novice agreements with expert scores for the EF sleep scoring method. The bars represent the mean ± standard deviation. The circles represent the specific results from each of the 8 files. B) A correlation between the EEG/EMG and EF 3-state sleep scoring results for each of the 8 files. Percent time spent in wake (blue circle), non-rapid eye movement (non-REM) sleep (green square), and REM sleep (orange triangle) were calculated for each file and represent the average score from the 3 expert scorers. C) Bar (mean ± standard deviation) and raw data for each recording representing the number of calculated state transitions for each scoring method (EF vs EEG/EMG) and scorer skill (novice vs expert). D) Novice #1’s 3-state scoring results for each file plotted against the average 3 expert scorers’ results for the EF sensor method. Novice #1 self-trained using only the Supplement 1 document. E) Novice #2’s 3-state scoring results for each file plotted against the average 3 expert scorers’ results for the EF method. Novice #2 used both Supplement 1 and Supplement 2 to iteratively self-train and improve sleep skill prior to sleep scoring data.

The combined 3-state scores for EF and EEG/EMG methods were highly correlated (R^2^=0.98) including when assessed within individual states (Figure 4B; R^2^>0.92). The number of state transitions into non-REM and REM sleep were not different between the two scoring methods, though EF results identified fewer transitions into wake than EEG/EMG (p=0.001) that were attributable to greater EEG sensitivity for brief arousals (Figure 4C).

Intra-scorer agreement between EEG/EMG and EF methods averaged 94.1 ± 0.5% with a single measure ICC of 0.90 ± 0.01 (Table 2) and aggregate sensitivity and specificity above 93% (Table 3). Of the 3 arousal states, REM sleep reported the lowest sensitivity and specificity for all comparisons, again as is typically noted given some ambiguity of the non-REM to REM transition period.

**Table 3.**
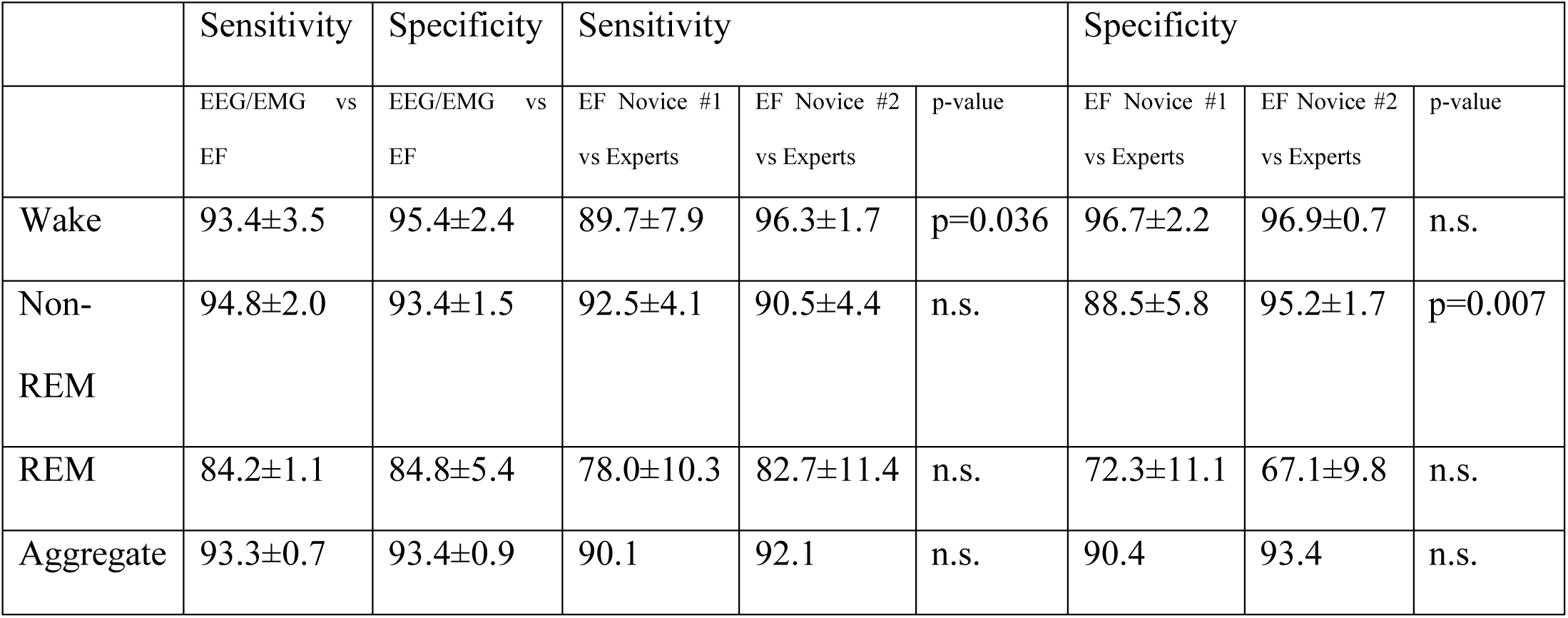
Summary of sensitivity (false positive rate) and specificity (false negative rate) when comparing the electric field (EF) sensor method against the electroencephalogram (EEG) and electromyogram (EMG) method (considered the ground truth) for scoring sleep. Novice scorer sensitivity and specificity, in which expert scorers are considered the ground truth, are calculated for the EF scoring method. Data are broken down by arousal state and summarized with an aggregate. P-values represent the comparison Novice #1 vs. Novice #2. Data represented as mean ± standard deviation. “n.s.” indicates no significance.

|  | Sensitivity | Specificity | Sensitivity |  |  | Specificity |  |  |
| --- | --- | --- | --- | --- | --- | --- | --- | --- |
|  | EEG/EMG vs<br>EF | EEG/EMG vs<br>EF | EF Novice #1<br>vs Experts | EF Novice #2<br>vs Experts | p-value | EF Novice #1<br>vs Experts | EF Novice #2<br>vs Experts | p-value |
| Wake | 93.4±3.5 | 95.4±2.4 | 89.7±7.9 | 96.3±1.7 | p=0.036 | 96.7±2.2 | 96.9±0.7 | n.s. |
| Non-REM | 94.8±2.0 | 93.4±1.5 | 92.5±4.1 | 90.5±4.4 | n.s. | 88.5±5.8 | 95.2±1.7 | p=0.007 |
| REM | 84.2±1.1 | 84.8±5.4 | 78.0±10.3 | 82.7±11.4 | n.s. | 72.3±11.1 | 67.1±9.8 | n.s. |
| Aggregate | 93.3±0.7 | 93.4±0.9 | 90.1 | 92.1 | n.s. | 90.4 | 93.4 | n.s. |

Reproducibility for both methods was assessed by blind inclusion of previously scored recordings. For both EEG/EMG and EF methods, average agreement of repeated recordings was above 96% with ICCs above 0.94 (Table 2), again in agreement with prior reports.^27^

### There was high EF scoring agreement between Novice and Expert

To determine the level of instruction needed to achieve high scoring agreement with experts, novice scorers were given an instruction document (Novice #1) as well as two sleep scored training files (Novice #2) to classify 3-state sleep scores with EF sensor data alone.

On average, Novice #1 produced agreements with the 3 expert scorers of 87.6±1.5 and an average ICC of 0.89±0.03 (Table 2). Overall sleep state identification for Novice #1 correlated well with the expert scorers (R^2^=0.96, p<0.001; Figure 4D). However, when broken down by state, wake and non-REM sleep scores correlated well (R^2^>0.86, p<0.001) but REM sleep scores did not (R^2^=0.004). Moreover, the number of state transitions into non-REM and REM sleep were lower (p=0.014 and p=0.006, respectively) compared to the expert scorers, but the number of wake transitions were not different.

Novice #2 produced higher percentage agreement with experts than Novice #1 (91.2±1.8, p=0.018) but had similar ICCs (0.91±0.02; Table 2). Novice #2’s overall wake and non-REM sleep state identification correlated well with the expert scorers (R^2^>0.97, p<0.001; Figure 4E). Importantly, Novice #2 REM sleep state scores were well correlated with expert scores (R^2^=0.69 p<0.001) unlike Novice #1 (R^2^=0.004 p=0.89). Novice #2 also produced higher wake sensitivity (p=0.036) and Non-REM specificity (p=0.007) to expert scorers than Novice #1 (Table 3).

### EF Sensors are able to Assess Sleep from Traditional Mouse Home Cages Without EEG/EMG

EF sensors placed on traditional mouse home cages were able to detect animal movement with high fidelity (Figure 3B). The voltage traces and spectrograms from these recordings appear indistinguishable from those collected during the validation experiment using the cylindrical acrylic chambers with EEG/EMG equipment. Moreover, 3-state sleep scoring was able to be performed from the EF data collected from the traditional home cages (Figure 3C_1_). The animals exhibited a biphasic sleep pattern during the dark cycle (Figure 3C_2_). Over the course of the 12-hour dark night cycle from 6pm-6am, the animals spent 17.8±5.6% of the night asleep (Figure 3C_3_). As expected, both the average time spent sleeping and the biphasic distribution of sleep match previously reported behaviors for the C57BL/6 strain of mouse.^28,29^

## Discussion

Knowing that respiration changes associate with sleep state,^5,8,19,22^ this work demonstrated that EF sensors could capture these changes to quantify 3-state sleep architecture with comparable accuracy as the gold standard EEG/EMG method. The EF sensor-based sleep scores had comparable or better agreement (i.e. less error),^7,8,12–14^ sensitivity (i.e. false positive rate),^7,12,13^ and specificity (i.e. false negative rate)^12,13,30^ than other studies comparing respiration-related changes to EEG/EMG to assess 3-state sleep architecture in rodents.

The EF sensor recording approach offers several advantages over other recording technologies. First, the approach is non-invasive and can be conducted on many mice more easily and cheaply than with a tether or surgically implanted sensor system. Furthermore, it allows recordings to be undertaken in the home-cage of pair-housed animals. It is worth noting the cages in this study were temporarily divided for pair-housed animals with an electrically shielded barrier to isolate animal recordings, however alternate strategies for isolating recordings from multiple animals within a single cage are possible. Dividing a home cage can limit social contacts to sight, smell, sound, and thermal exchange during recording sessions and should be realized as an important ethological factor in future experimental designs. Other advantages of the EF sensor approach include that; 1) sensors can be placed outside the animal’s environment, 2) can be undertaken in both rats and mice with equal ease,^23^ 3) there is no required specialized software for analyses, 4) components are inexpensive to purchase and easy to construct,^23^ and 5) remove the need to used cranial electrodes to capture sleep thus freeing them to record other EEG features.

Importantly, EF sensor recordings also have the potential to simultaneously quantify other motor behaviors such as respiration profiles in non-REM and REM sleep, grooming, locomotion, eating, and drinking using only a frequency-based features of a single voltage channel.^23^ Sleep scoring using the EF sensor data was also easy to learn and novices were able to achieve comparable results as expert sleep scorers. In the future, scoring based on EF voltage traces and spectrograms provides rules that may form the basis for automated detection algorithms that may improve sleep analyses.

EF sensor sleep scoring has inherent limitations common to most sleep scoring methods. As with EEG/EMG scoring, the majority of error occurred at state transitions, brief arousals, and at the non-REM to REM transition.^31,32^ EF sensors detect only movement-related variables, and brief electrocortical arousal events that are visible on EEG can occur independent of movement. Consequently, the developed rules that rely on detection of movement-related variables required longer periods than EEG/EMG for the EF sensor method to be scored as wake. This led to missed brief arousals and likely contributed toward less accurate assessment of brief arousals, seen as reduced transitions into wake (Figure 4). Since brief arousals account for a small percentage of total assessed time, missing these brief events does not affect overall wake time but could negatively impact calculation of other sleep measures (e.g. sleep event duration or sleep fragmentation).

Similar to missed brief arousal detection, transitions into REM sleep were another common source of error for EF sensor sleep scoring. The EF sensors, again, detect movement but REM sleep may show up on an EEG slightly prior to the start of summed body movements detected by the EF sensors. Often this discrepancy was a single 10-second epoch, but occasionally more. REM-related scoring error is also common in EEG/EMG sleep scoring methods partially due to their low occurrence (typically 5-15% of total sleep time)^31,32^ and placement of EEG electrodes.^33^

Overall, novice scorer error was better than reported values for other respiration-related, non-contact methods to assess rodent sleep.^8^ Accurately quantifying REM sleep events proved to be the poorest agreement with the expert scorers due largely to Novice #1 missing REM events entirely. Novice #2 was able to improve on Novice #1’s REM sleep scoring challenges putatively because Novice #2 also received samples of EF scored sleep (Supplemental File 2). These sleep-scored samples allowed Novice #2 to score recordings then compare the results to the expert scores provided. This process increased accuracy of sleep scoring by correcting mis-classification. Novice #2’s overall agreement with experts was nearly similar to the inter-expert scoring variability, but, as with Novice #1, REM events were the source of greatest error. This suggests that accurate determination of REM sleep to be the most challenging aspect of sleep scoring for the EF method (and arguably EEG/EMG methods). This is worth noting for other groups wishing to implement EF sensor sleep assessment into their work. Overall, with appropriate training resources provided, researchers with no sleep scoring experience may be able to undertake sleep scoring with comparable accuracy to experts using traditional EEG/EMG methodology.

EF sensors were also able to continuously capture 3-state sleep in traditional mouse home cages overnight, without the need for EEG/EMG equipment, and reproduced previously reported sleep patterns for same C57BL/6 strain of mice.^28,29^ Though only overnight data was presented in this study, the EF sensors have since proven to be easy to use, consistently and continuously measure animal movement over days and weeks, as well as detect behavioral differences between healthy and injured mice of multiple models and strains (unpublished observations). These strengths are tempered by environmental limitations that affect the EF sensors recording quality,^23^ and are important to emphasize here. First, response magnitude is highly distance – dependent indicating magnitude of voltage response may not itself identify a movement related event. In related fashion, placement of additional sensors at multiple locations may be required to accurately detect smaller movements including respiration. Second, EF sensors are dependent on the electrostatics of the environment which are reduced with humidity.

Overall, the EF sensors scored sleep accurately and novice scorers were able to reproduce expert sleep scores using only self-training. The EF sensors are adaptable, noninvasive, and able to detect a wealth of behavioral information continuously over long timelines. Though the EF sensors have limitations, they provide a more ethological method to assess sleep in the animal’s home cage without need for surgical implants, including in cage environments unsuitable for EEG/EMG equipment.

## Acknowledgments

This work was supported by grants from the Craig H. Nielsen Foundation (to SH) and the NIH (K08NS105929 to NPP and 5k12-Gm000680 to HK).

## Abbreviations

EEG: electroencephalogram
EMG: electromyogram
EF sensors: electric field sensors
REM sleep: rapid eye movement sleep
Non-REM sleep: non-rapid eye movement sleep

## Disclosure Statement

### Financial Disclosure

HK, WG, and SH are co-inventors of US patent application 16/095,906, filed 10/23/2018, that includes use of EF sensor methodology for non-contact physio-behavioral monitoring of movements including respiration.

