## Supplement 1 for "Noninvasive Sleep Scoring in Mice using Electric Field Sensors"

### Criteria for Scoring Sleep Using Electric Field Sensors

### Contents

- This document will help you understand how to score sleep using electric field (EF) sensors.
- Sleep scoring rules are on slide 14 of this document
- Discussion topics include:
  - Sleep stages
  - What does each arousal stage look like on the EF sensors
  - Rules for scoring sleep using EF sensors
  - Pitfalls and troubleshooting

### Possible Stages:

- Wake
    - Characterized by awareness and responsiveness to environment
    - Includes the quiet rest time before you fall asleep
- Sleep
    - Rapid Eye Movement (REM) Sleep
        - Paralysis of non-respiratory muscles (except for brief twitches and eye movement)
        - Enriched dream content; irregular and more shallow respiration
        - Makes up ~ 5-10% of your total sleep time
        - In normal sleep, occurs after Non-REM and often followed by a brief arousals
    - Non-Rapid Eye Movement (Non-REM) Sleep
        - Decreased responsiveness, regular respiration, some movement
        - Rodents Non-REM is not further divided into the subcategories that exist in human Non-REM sleep
        - Makes up the remaining 90-95% of total sleep time

### Electric Field (EF) Sensors and Sleep

- Raw Voltage Trace
  - The EF sensors translate movement into a voltage trace
  - The larger the movement, the bigger the voltage signal. Different animal movements will show up as unique voltage patterns (on next slide)
  - When the animal is sleeping, it is still moving (breathing, twitches during REM)
- Spectrogram
  - The different patterns in the voltage trace are easier to see when you convert them into a spectrogram
  - A spectrogram visualizes the frequencies and relative representation (power) of each frequency in a voltage trace

**Raw Voltage Trace:**

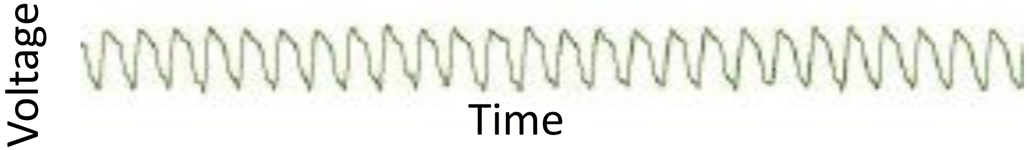


**Spectrogram:**

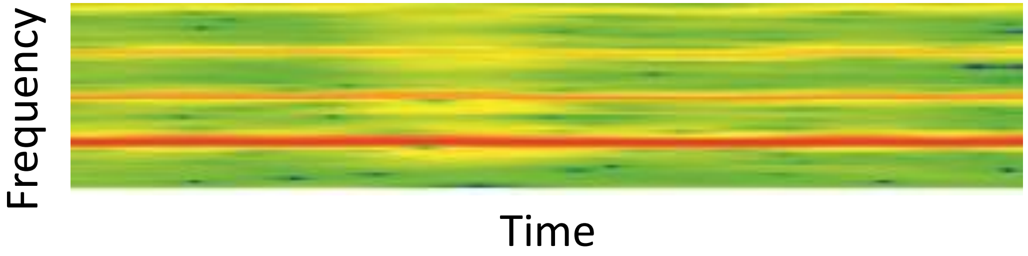


Color denotes the relative strength (power) of the frequency. Red means high power (i.e. that frequency is strongly present), green/blue means low power (i.e. that frequency has weak or no presence)

### What does WAKE look like on the EF sensors?

Exploration

**Raw Voltage Trace:**

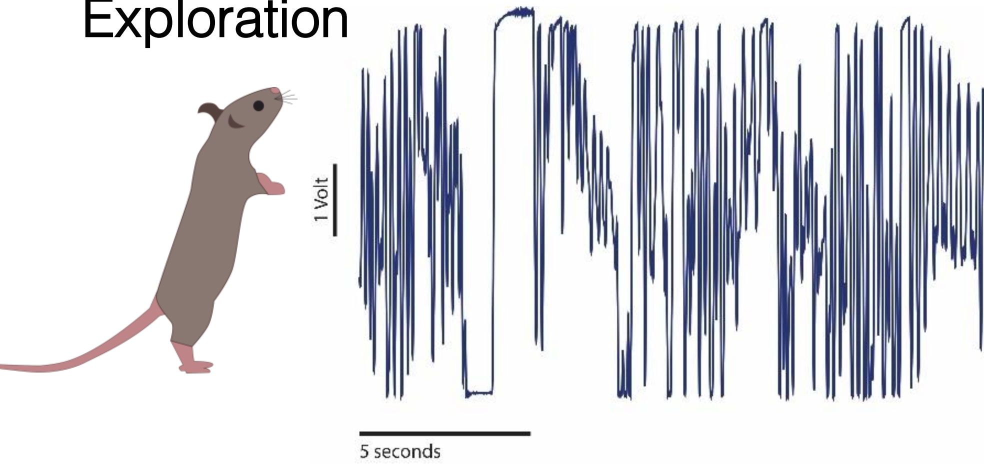


Grooming

**Raw Voltage Trace:**

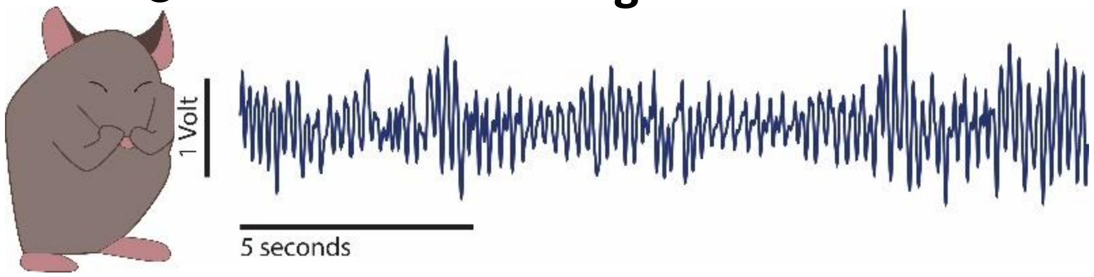


Eating

**Raw Voltage Trace:**

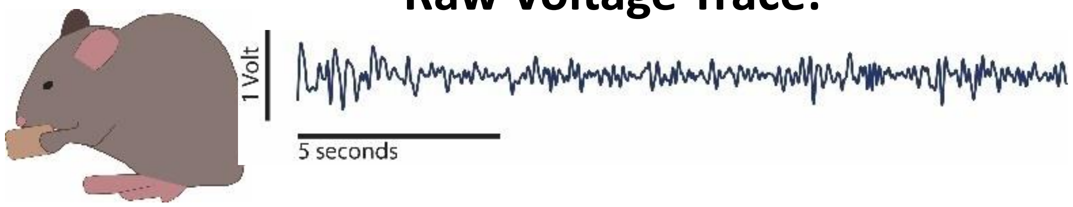


Drinking

**Raw Voltage Trace:**

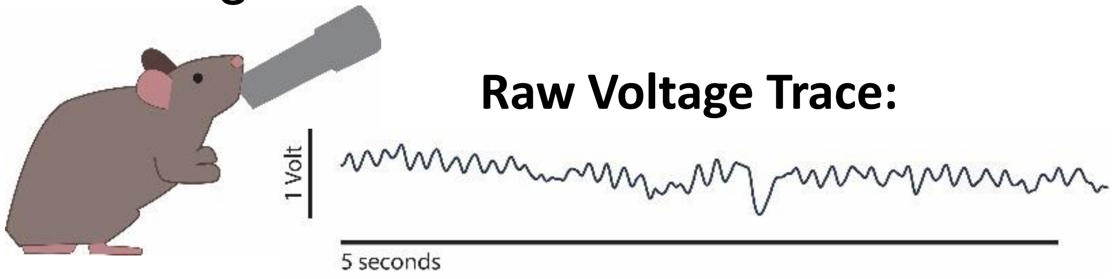


- EF sensors record Wake as many different voltage patterns

### More WAKE Examples with Spectrograms

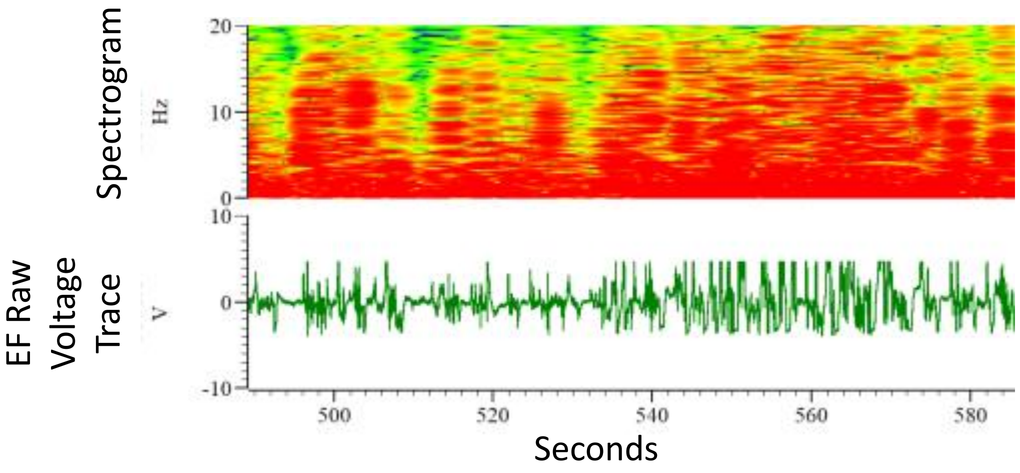


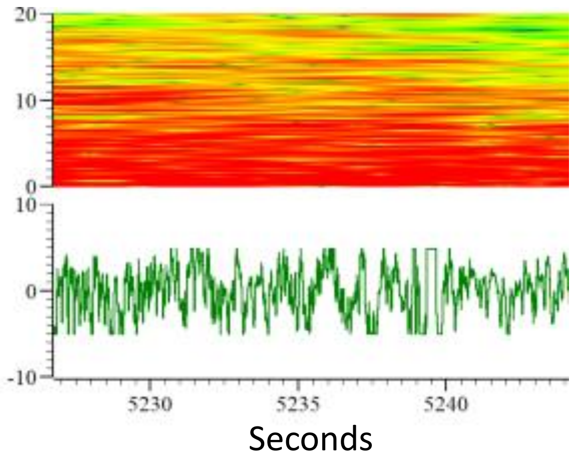


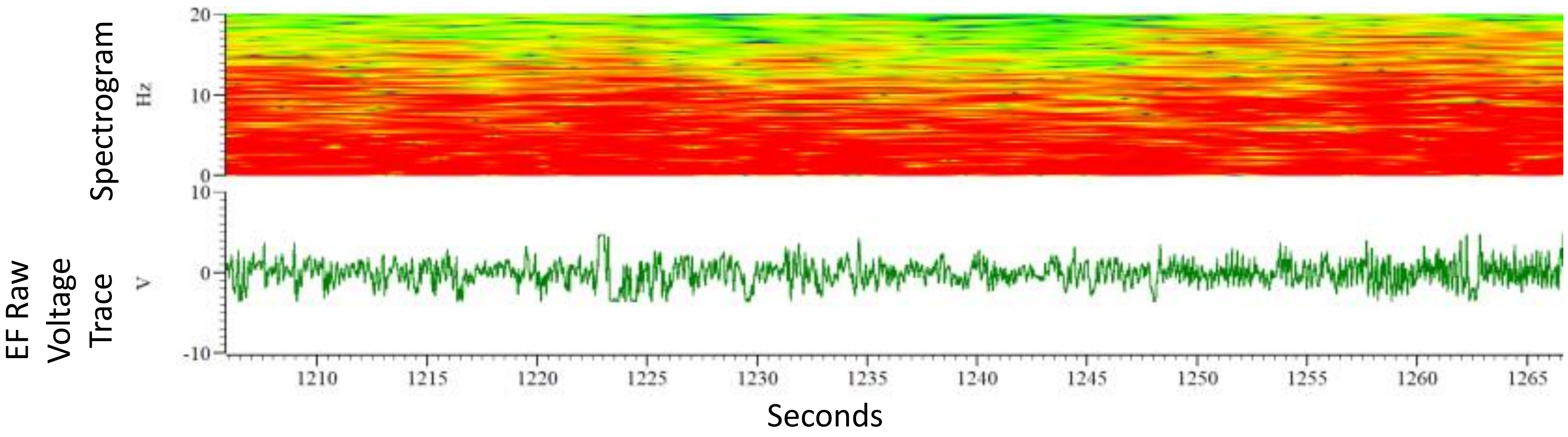


- The animal is typically moving or sniffing while awake.
- Definition of WAKE:
  - Raw Voltage Trace
    - **Large** amplitude (relative)
    - **Erratic** shape with **high variability** of amplitude and width
  - Spectrogram
    - Powerful (i.e. more red) signal
    - Solidly covers frequencies ~0 to 15-20 Hz (there is a caveat to this because grooming, eating, and drinking sometimes show up with a slightly stronger wide and messy band between 5-10 Hz)

### What is Non-REM Sleep for the EF Sensors?

**Examples of Voltage Trace Patterns of Respiration During NREM Sleep**

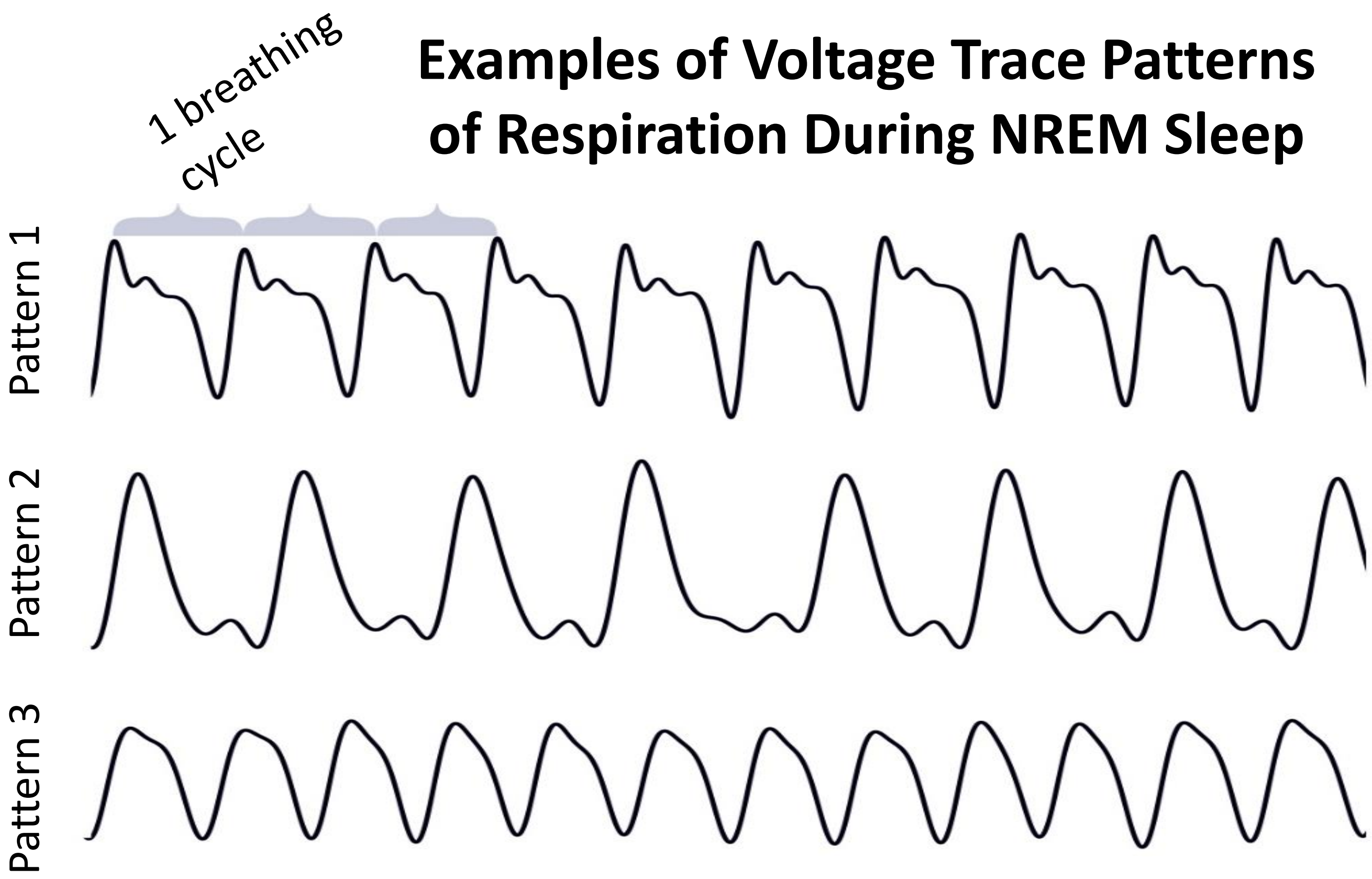


- During Non-REM sleep, the animal is still moving because breathing expands and contracts the chest wall (this shows up as a consistent cyclic pattern on the EF sensors).
- The cyclic pattern will change based on the animal's orientation to the sensors (see left), but the shapes will remain consistent during a Non-REM event.

### Non-REM Sleep Examples

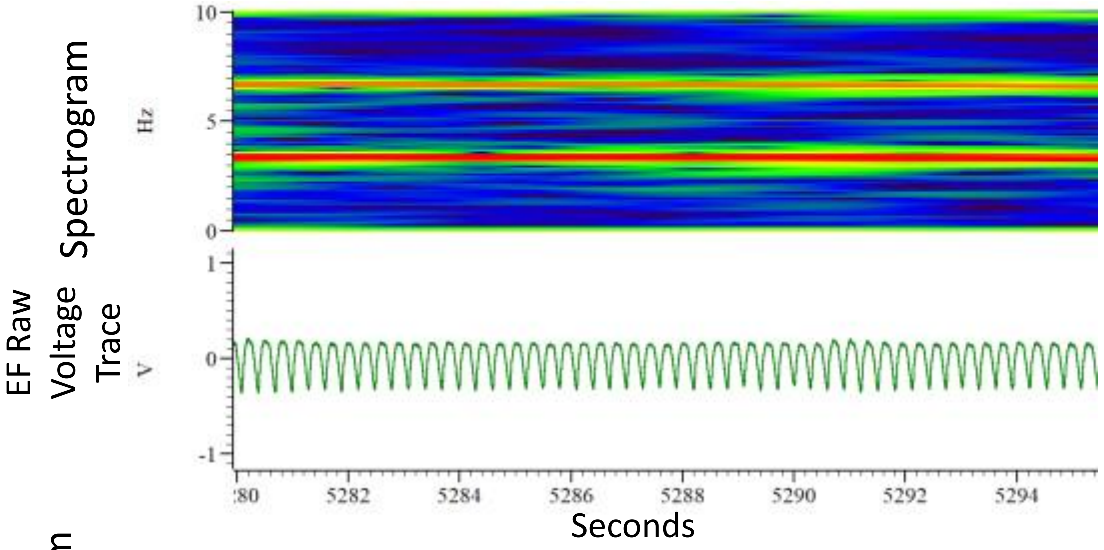


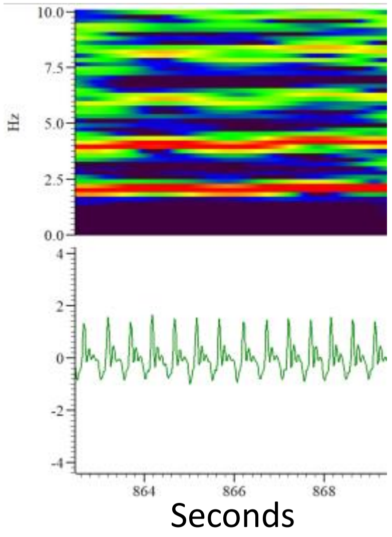


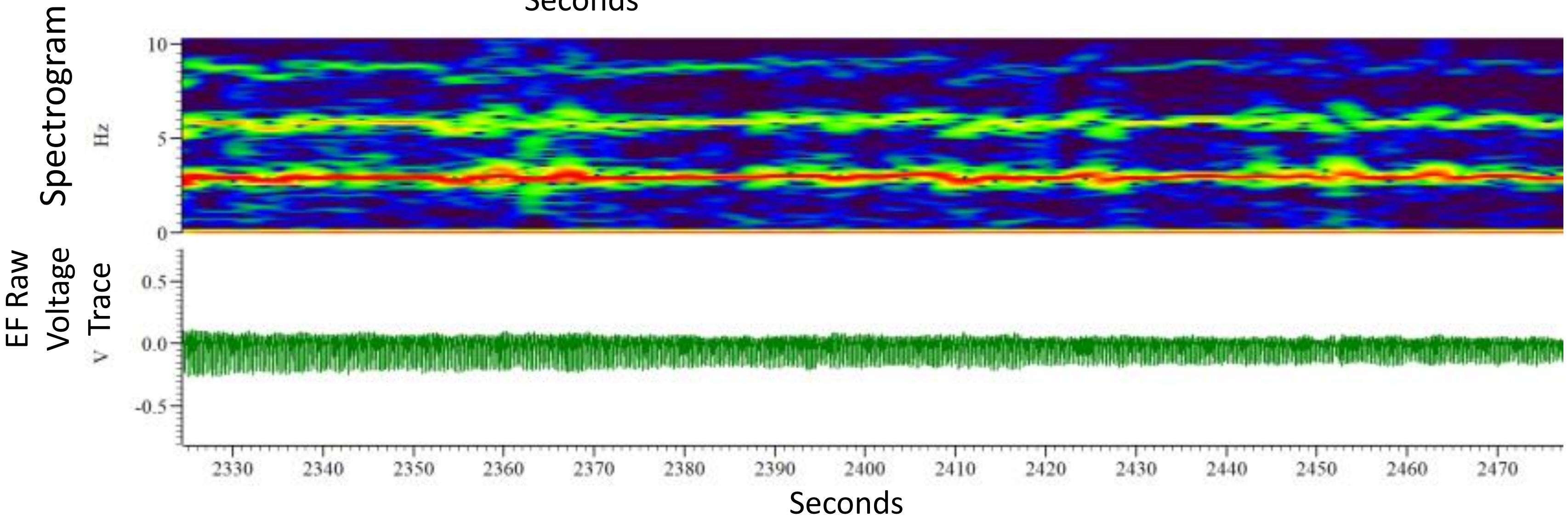


- The animal typically stays in one place during Non-REM sleep.
- Definition of NREM Sleep:
  - Raw Voltage Trace
    - **Low** amplitude (relative)
    - **Consistent** shape with **consistent** amplitude and width
  - Spectrogram
    - Powerful (i.e. more red) single band between 2-4 Hz against a relatively less powerful (i.e. more blue and green) field.
    - Very little to no other frequencies present
    - May have less powerful harmonic lines at frequency multiples of band (this is because the waveform is not a perfect sine wave).

### What is REM Sleep to the EF Sensors?

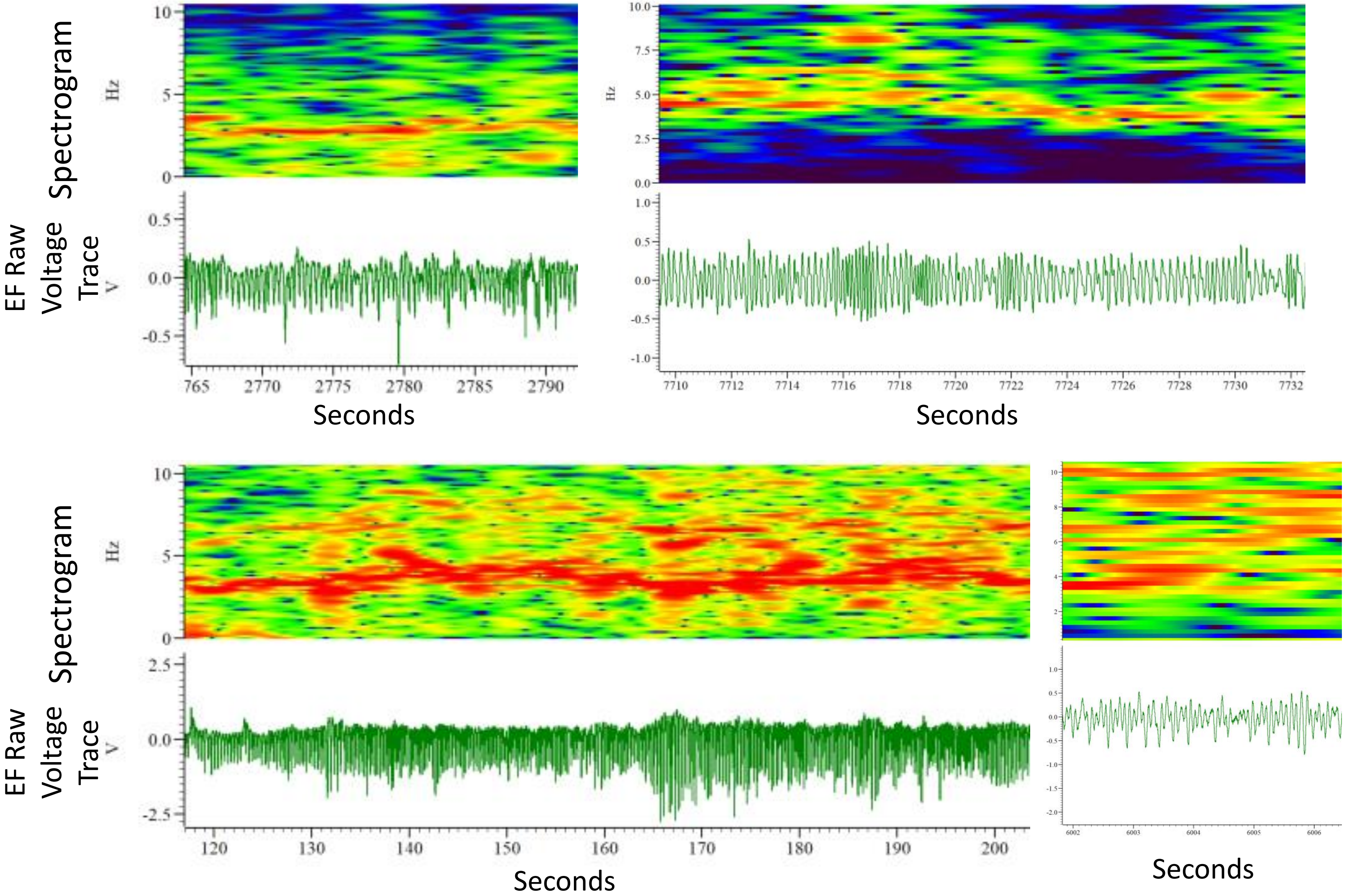


- During REM sleep, there are small twitches of fingers, ears, eyes, and toes, but breathing is still the dominant motion.
- These twitches show up on the voltage trace and “muddy” the cyclic breathing pattern.

### REM Examples

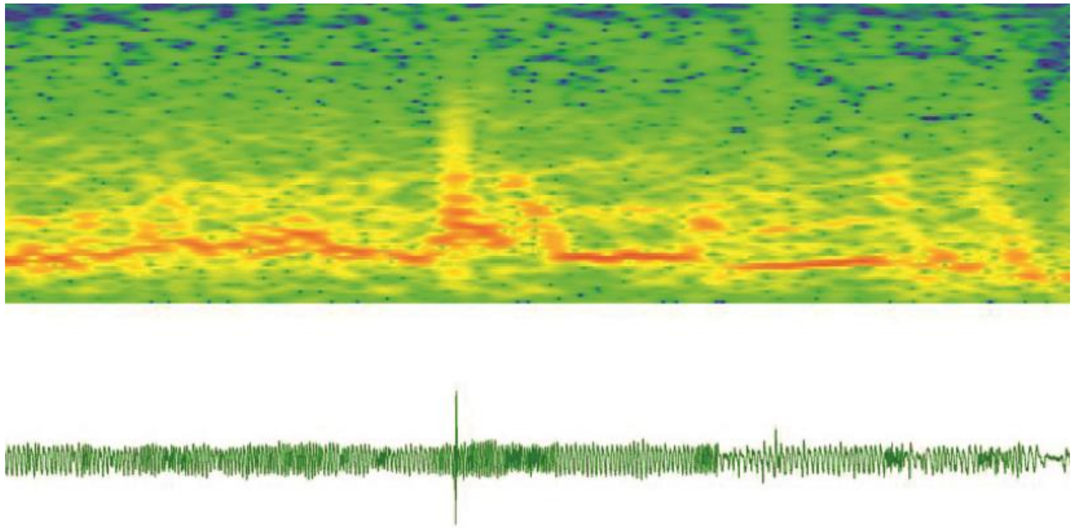

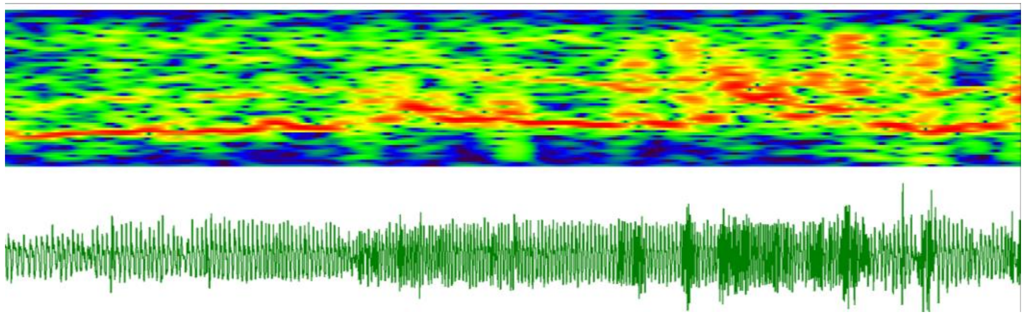

This example is entirely REM, but there are two distinct patterns – that's ok!

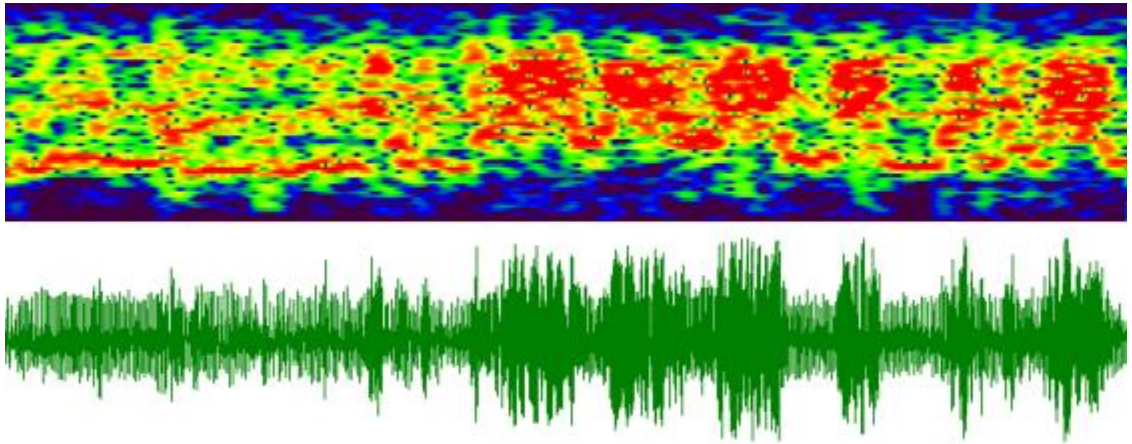

- The animal is mostly still during REM sleep.
- Definition of REM Sleep:
  - Raw Voltage Trace
    - **Similar** amplitude to preceding Non-REM signal
    - **Somewhat Erratic** shape (relative to Non-REM signal) with **Erratic** sinusoidal amplitude and width (but not so much as WAKE)
  - Spectrogram
    - Fragmented, spotty red pattern between 1-10 Hz (there should be NO meaningful signal <1 Hz, otherwise it might be WAKE).

### Putting All the Stages Together – getting a sense of transitions

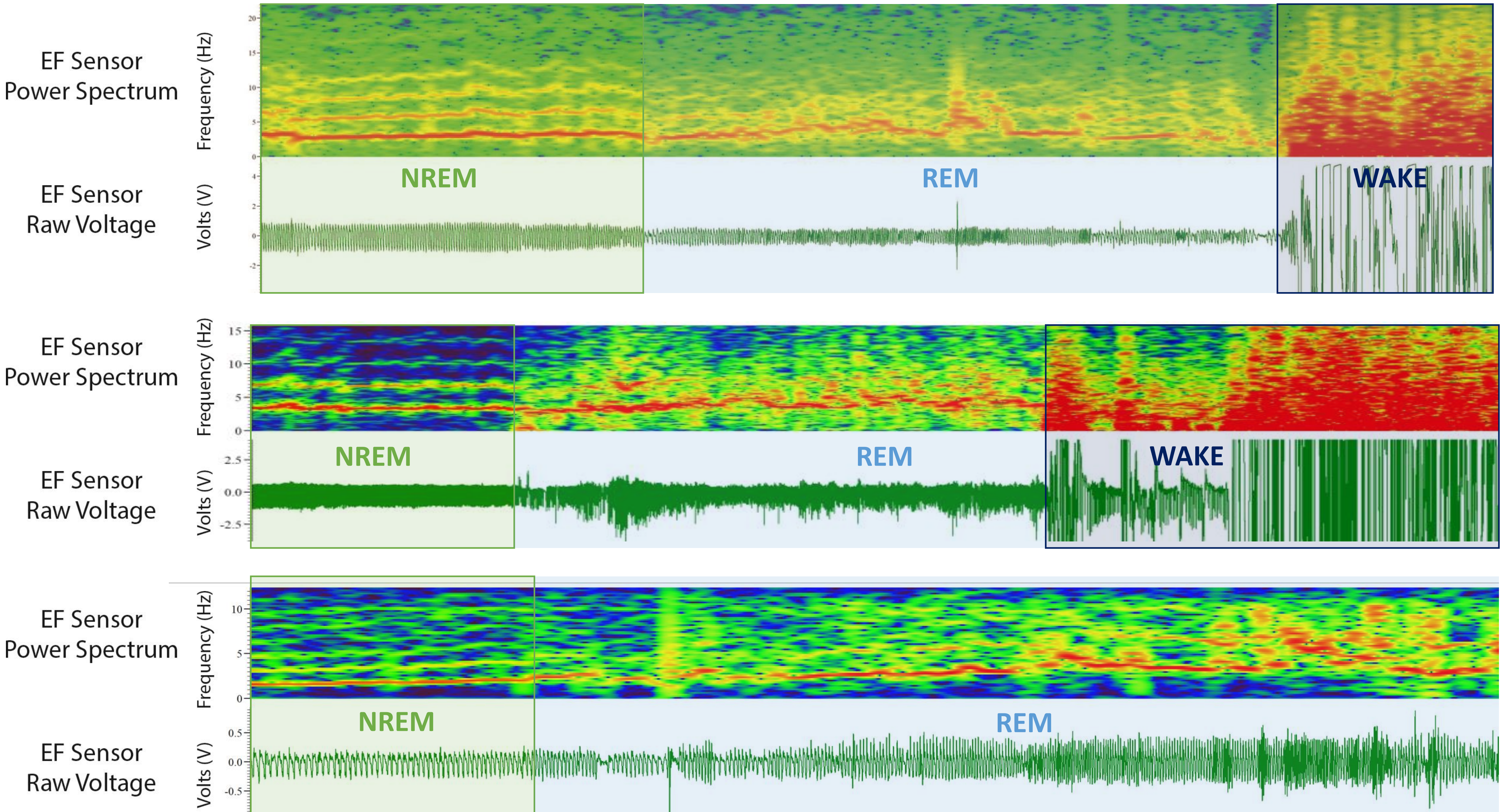

### More Examples…

**This portion is not long enough to count as an arousal (WAKE) – we'll talk about how to label this later**

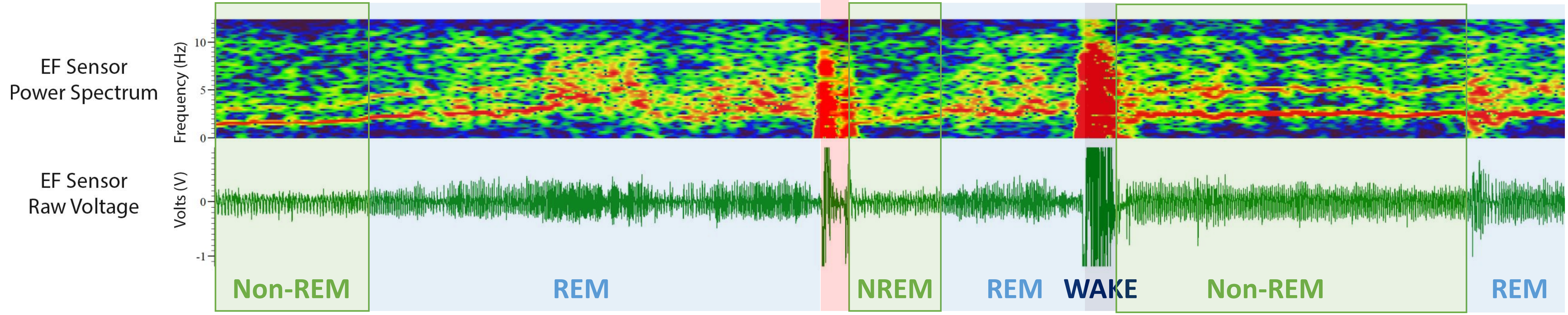


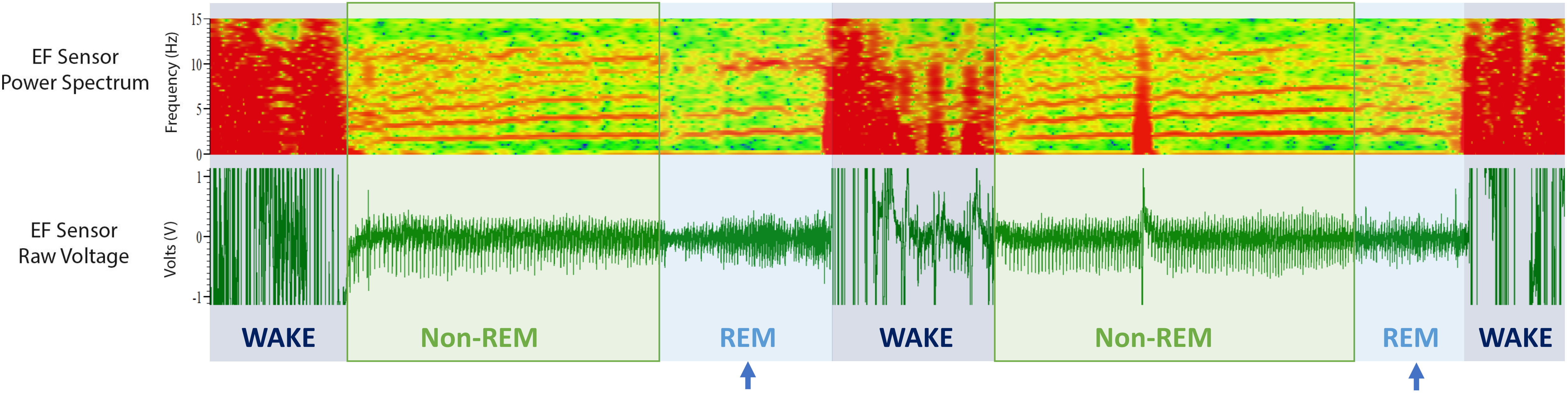


**These REM portions are subtle with strong respiration signal still – you have to look at the raw voltage trace and the >5Hz frequency area**

### How is Sleep Scored

- Sleep is scored by dividing your data into individual chunks (called epochs) of a few seconds and assigning a state (WAKE: W, Non-REM: N, or REM: R) to each epoch:

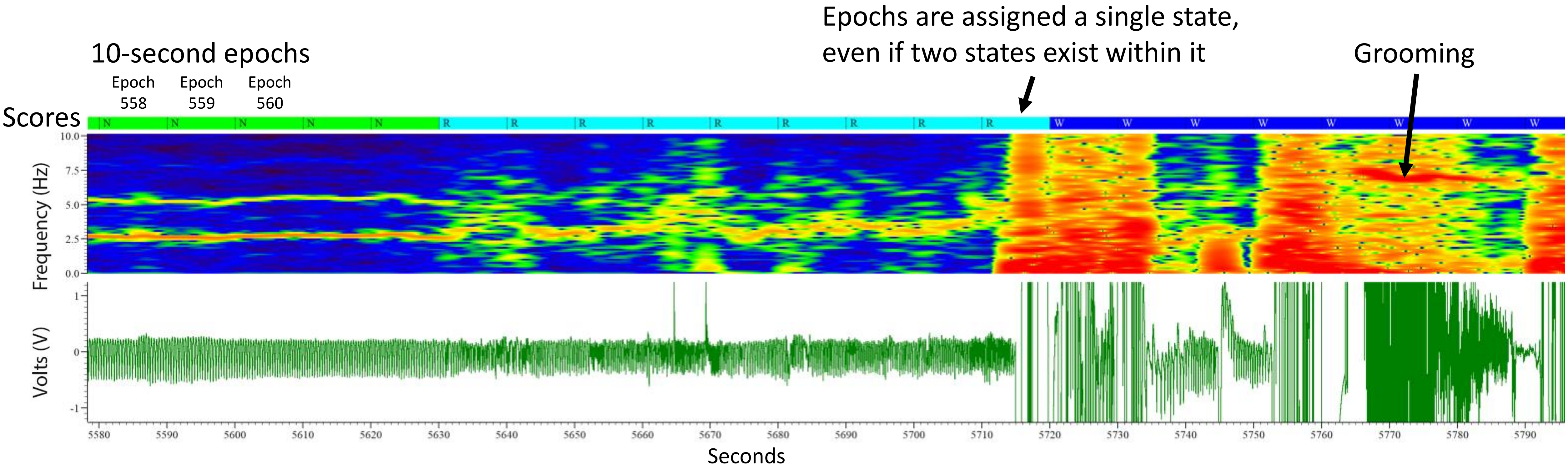

### Rules for Scoring Sleep in Mice Using EF

- Each epoch is scored based on the majority of the signal present in that epoch
  - i.e. if a 10-second epoch has 7 seconds of WAKE and 3 seconds of Non-REM, it will be scored as WAKE
  - If an epoch is evenly divided b/t two states, score it as the immediately previous state
- REM **CANNOT** transition directly from WAKE without Non-REM between
- Non-REM **CANNOT** exist as a single epoch
  - You must have two+ consecutive epochs of Non-REM to change the state to Non-REM
  - You **CAN** have a single epoch of WAKE, but only if it is 10+ seconds long **AND** follows the majority rule above

### Troubleshooting and More Examples

- More information about handling brief arousals
- More examples of scoring transitions

### When you have a single WAKE epoch in a sea of Non-REM

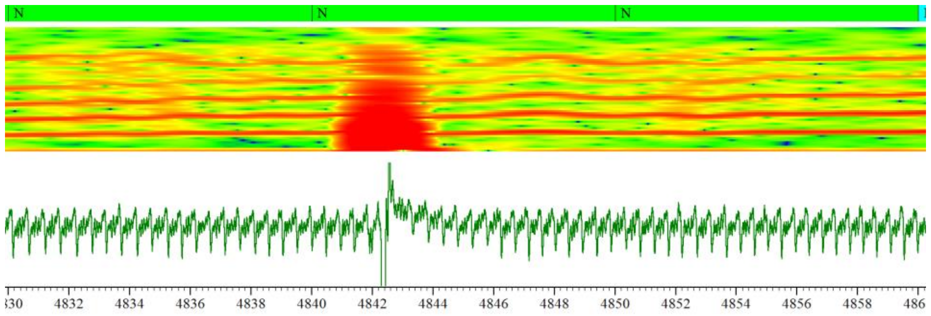

- This is a twitch, the actual movement on the voltage trace only lasts ~3 seconds, so there is no scored WAKE state change

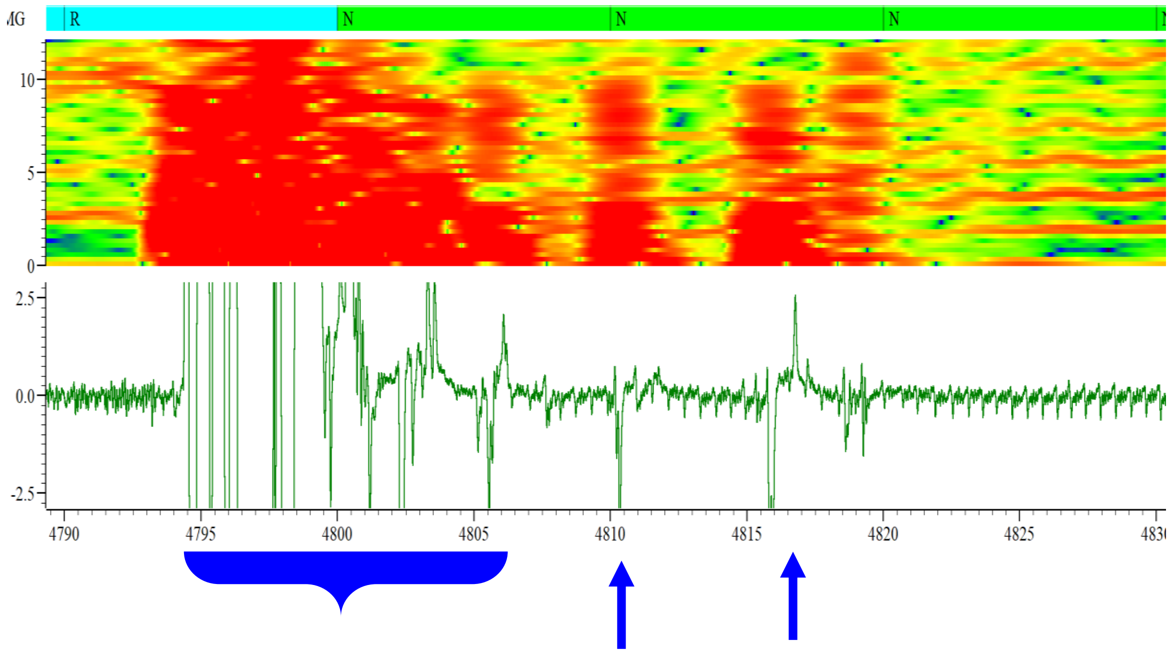

- The WAKE behavior (bracketed left) signal meets the 10+ second criteria, but not the majority rule b/c it straddles two epochs without taking a clear majority for either
- The blue arrows are independent twitches that are not part of the previous 10-sec movement. This is because you can see distinct and even respiration cycles between the twitches

### More Examples

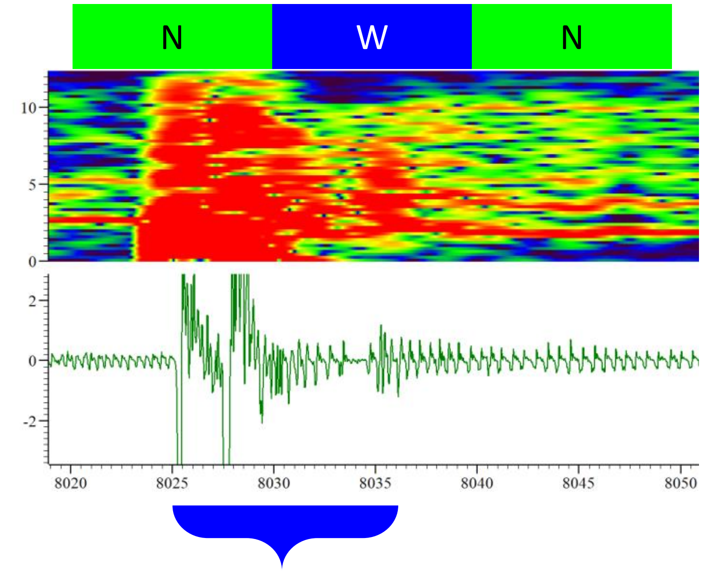


- This event meets the 10+sec and majority rules (though it BARELY meets the majority rule)
- The first epoch is still scored as Non-REM b/c WAKE does not occupy a majority of the epoch so it is scored as the previous state.

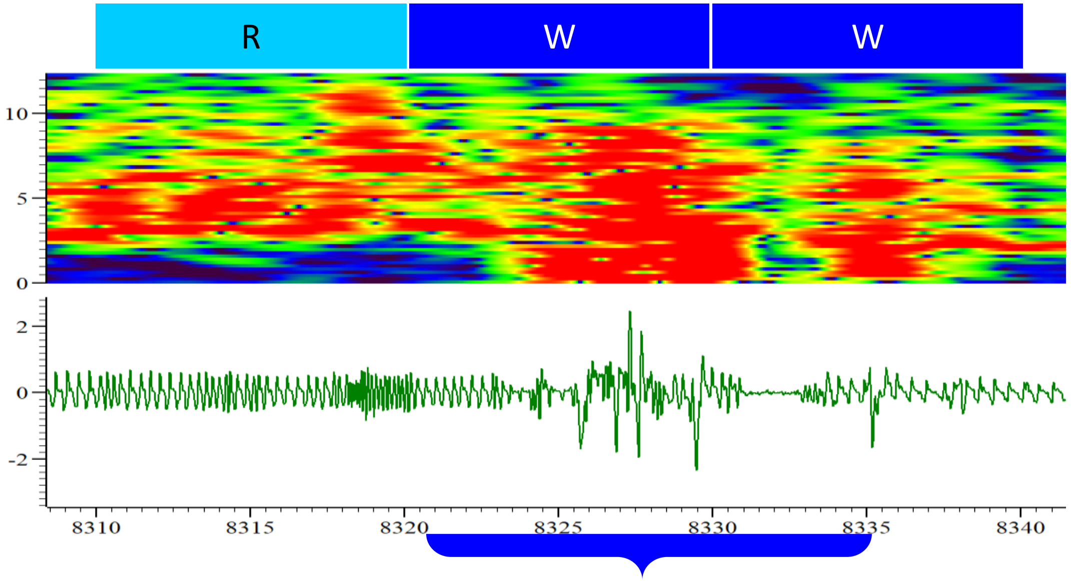


- This event meets the 10+sec and majority rules
- The middle epoch is scored as WAKE b/c REM does not occupy a majority of the epoch.

### Remember this?

**This portion is not long enough to count as an arousal (WAKE) – we'll talk about this now ☺**

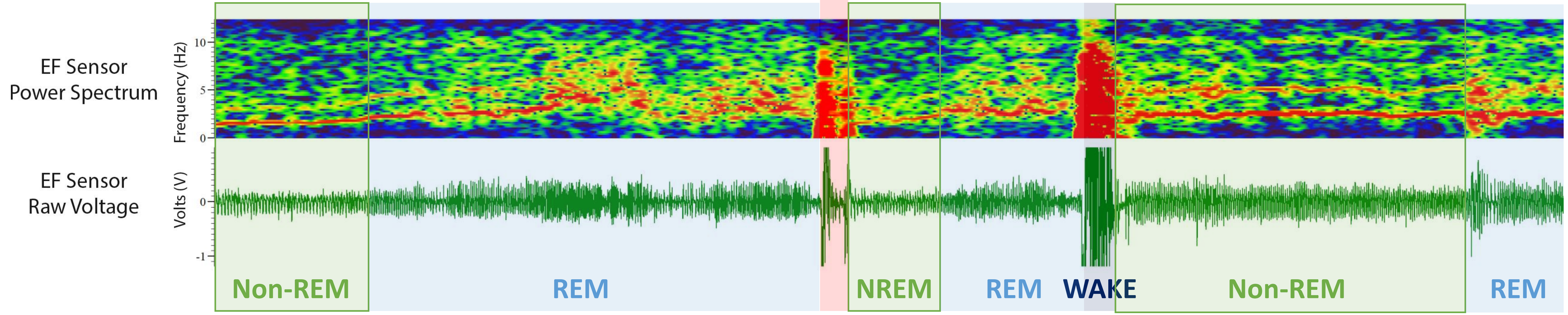


This portion gets scored as Non-REM b/c the arousal does not meet the 10+ sec or majority criteria. This does not get scored as more REM b/c REM ends with the arousal movement

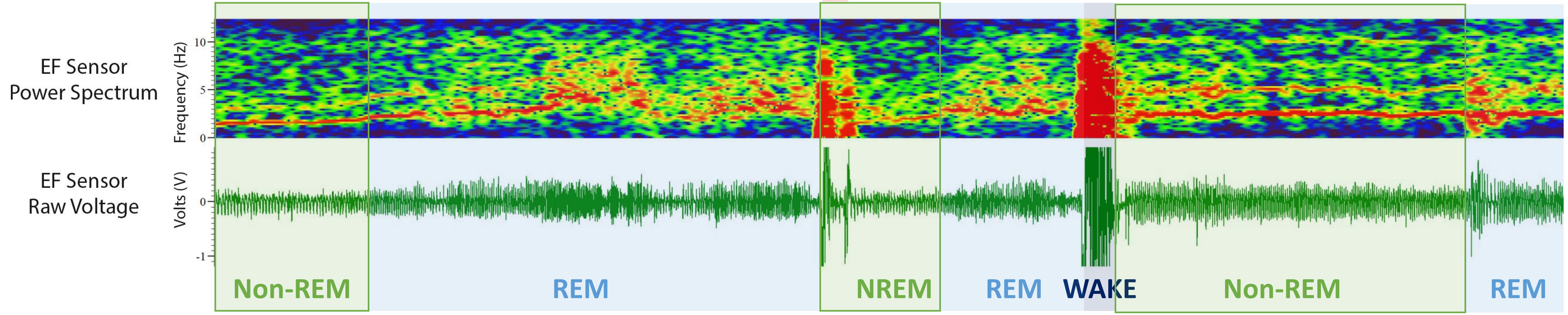
